# Transposon activities drive the selection and diversification of sweet orange (Citrus × sinensis) cultivars in the last 500 years

**DOI:** 10.1101/2022.03.19.484946

**Authors:** Bo Wu, Yiping Cui, Yongping Duan, Frederick G. Gmitter, Feng Luo

## Abstract

Sweet orange (SWO) has one of the largest numbers of cultivar groups in *Citrus* with diverse horticultural traits just through asexual breeding. However, the molecular mechanism driving its fast selection and diversification is unclear. In this study, we completely surveyed the transposon activities in SWO genomes and unraveled six transposon families with up to 8,974.2-fold activity increase in modern SWO cultivars. Based on the transposon insertion phylogenetic tree, we inferred that modern SWOs date back to a common ancestor ∼500 years ago and reconstructed three major spread events in SWO cultivation history. Activation, acceleration, and silencing of the six families have recurred in distinct lineages. Their insertions are presented as tag mutations for all SWO cultivar groups and can distinguish over 99% of all analyzed SWO accessions. The insertions are enriched in impacting plant development and hormone signaling. This study demonstrated the importance of transposon activities in asexual breeding.

## Introduction

Inter-specific hybridizations between pummelo (*Citrus maxima*) and mandarin (*Citrus reticulata*) created the original sweet orange (*Citrus* × *sinensis*, SWO)^1, 2^, but when and where have yet to be discovered. Although the Chinese character ‘橙’ (orange) was mentioned as early as the 2^nd^ century BC^3^, it is usually referred to as sour orange (*Citrus × aurantium* L.) or Xiangcheng (*Citrus × junos*) in ancient Chinese literature rather than *C. sinensis*. Even in *Ben Cao Gang Mu*, a book completed in 1578 AD, Shi-Zhen Li described orange fruit as sour but mandarin fruit as sweet and sour, indicating oranges were most likely referred to as sour oranges then. However, by just asexually propagation through apomixis and grafting in several centuries, SWO has one of the largest numbers of bud sport selections in *Citrus* with diverse horticultural traits^2, 4–6^, reaching over 1,300 recorded accessions already in 1936^7^ and many more since then. The molecular mechanism driving this fast selection and diversification of SWO is not well understood. Although somatic mutations have been regarded as responsible for the phenotypic diversification of SWO cultivars, researchers have detected very few tag mutations of SWO cultivar groups before^4, 5^.

Many asexually propagated economically important horticultural cultivars have been observed with high transposition activity, which was proposed to be due to the relaxation of selective pressure^8^. Nevertheless, loose pressure only interprets the random activation of TEs in a few individuals but cannot explain the transposon activity increase of a whole species or population. On the other hand, asexually propagated horticultural plants are subjected to continuous artificial selection of bud sports (somatic mutants), phenotypically distinct parts of plants^9^. Artificial selections, resembling positive selections in nature, could have led to the widely observed higher transposition activities of asexually propagated horticultural plants instead^10^. The fact that distinct stresses have been reported to trigger transposon movement^11^ implies that TEs are potentially utilized as mutagens in adaptative evolution. Microbial experiments and simulations also showed that positive selections more easily favor individuals with higher mutation rates in asexual organisms^12, 13^. However, similar evolutionary experiments are difficult to carry out for horticultural plants with much longer generation time. SWO includes several popular cultivar groups with partially recorded breeding timelines, including Navel orange, Valencia orange, Blood orange, Jincheng, and Bingtangcheng. TE insertions have been reported to contribute to the formation of Europe-origin Blood oranges and the China-origin acid-less Bingtangcheng^2, 5^. These unique features of SWO^12^ make it a model horticultural species for studying the activity dynamics and contribution of TEs in the asexual breeding of horticultural species.

In this study, we developed a bioinformatics pipeline to completely survey the TE activities in SWO genomes and detected 34 active TE families in SWO. We unraveled their insertion atlas across major SWO cultivar groups with high precision and sensitivity. Statistical analyses led to the discovery of six TE families with significant transposition activity increase in modern SWO cultivars dating to a common ancestor ∼500 hundred years ago, while most of them have remained low activity levels in early diverging local lineages under far fewer artificial selections. The boosted TEs experienced multiple transposition activation, acceleration, and silencing events independently in distinct lineages. Their insertions are presented as tag mutations for all SWO cultivar groups and can distinguish over 99% of all analyzed SWO accessions and are enriched in impacting plant development and hormone signaling. Our study shows that TEs have played a crucial role in SWO breeding and their activity levels in horticultural species are closely related to artificial selections.

## Results

### High-accuracy allele-aware transposon insertion scanning in sweet orange

To detect active TEs, we compared eleven published SWO genomes, including Navel oranges, Valencia oranges, and three native cultivars from China (Figure 1a and Table S1). Interspersed repeats with at least two distinct insertion sites among the eleven assemblies were identified as putatively active TEs (see Methods). As a result, we found a total of 34 such TE families, including 11 hAT, 10 MULE, 5 Harbinger, and 1CACTA-type DNA transposons, 4 LTR (long terminal repeat, 3 Gypsy and 1 Copia)-type retrotransposons, and one of unknown type. Representative members of these TE families in the DVS assembly are listed in Table S2. To study the historical TE activities in SWO, we collected whole-genome next-generation sequencing (NGS) data from 127 SWO accessions covering all major sweet orange cultivar groups (Table S3).

**Figure 1.**
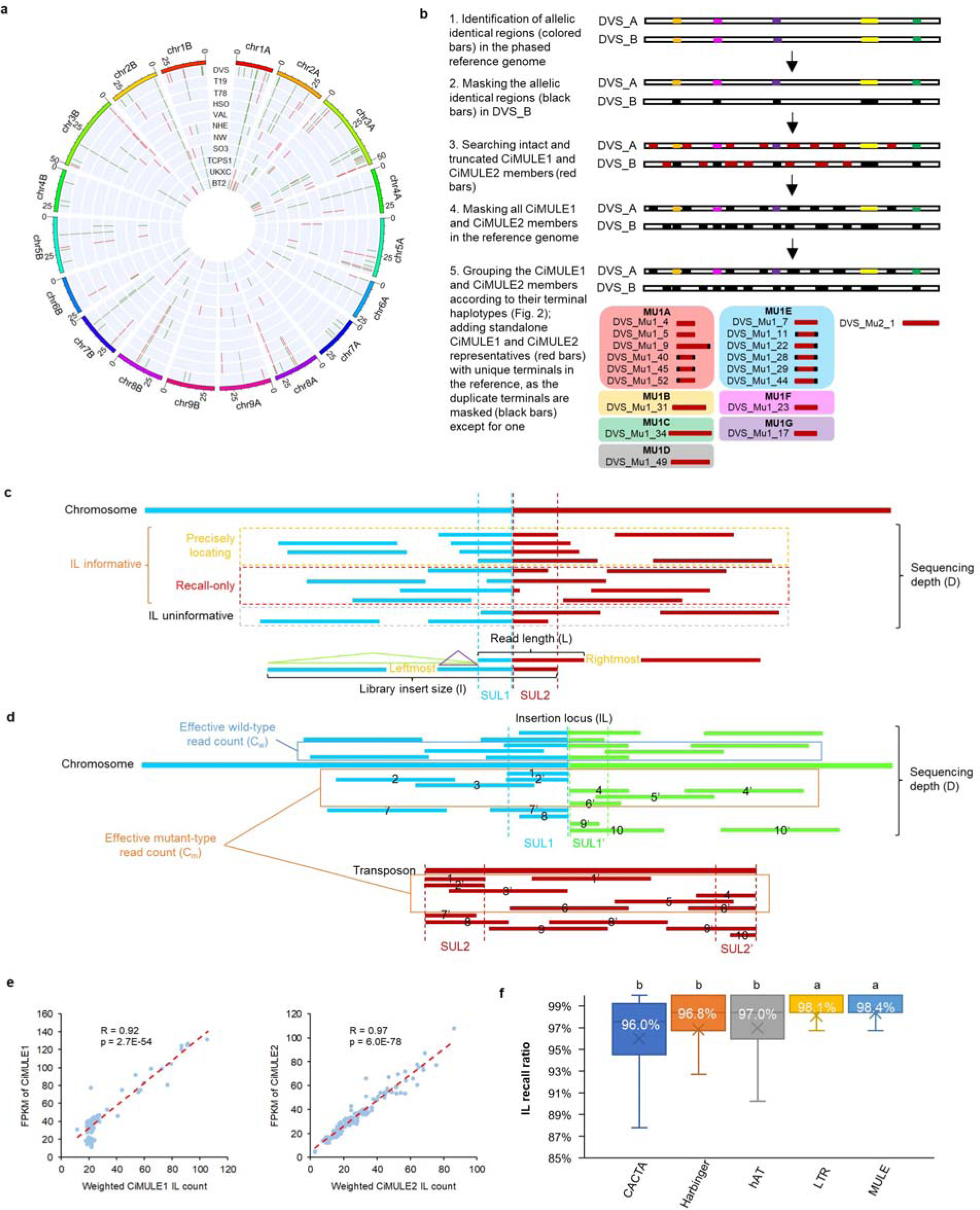
Detection of active transposon families and their insertions in sweet orange genomes. **a),** exemplary insertion atlas of active transposons in SWO assemblies. A total of eleven SWO assemblies including DVS and the diploid reference were compared. The reference (chr1-9A, B) is shown utmost, with the tick mark unit in million base pairs (Mb). Green and red bars denote CiMULE1 and CiMULE2 insertions, respectively. Aligned assemblies: T19 and T78, two irradiated Valencia sweet orange mutants; HSO, di-haploid Valencia sweet orange genome; VAL and SO3, diploid Valencia sweet orange assemblies; NHE and NW, navel oranges; TCPS1, UKXC, and BT2, sweet orange cultivars from South China. Sources of these assemblies are listed in Table S1. **b)**, modification of the diploid reference genome for high-quality transposon insertion locus (IL) scanning. CiMULE1 and CiMULE2 are used as examples. DVS_A and DVS_B are the two homologous chromosome sets of DVS. The bars with the same colors (except for the red) on DVS_A and DVS_B denote allelic identical regions, red bars denote the transposons, and black bars indicate masked regions. **c)**, mapping of pair-end sequencing reads to transposons embedded in the reference genome. The blue and green (in panel **d**) bars denote upstream and downstream regions/reads of transposons (red bars). IL overlapping read pairs are classified as informative and uninformative in recalling the IL. Precisely locating reads can be used to precisely locate and recall an IL recall, while recall-only read pairs could only recall an IL. The leftmost and rightmost read pairs allowing recalling of the IL are shown at the bottom. SUL (shortest unique k-mer length) denotes the shortest length of the corresponding region allowing unique mapping in the reference genome. **d)**, mapping of pair-end reads to novel ILs absent in the reference. Wild-type and mutant-type pair-end reads are drawn on top and below the chromosome, respectively. Mutant-type read pairs are either dis-concordantly mapped or include at least one read split-mapped to both the IL surrounding sequences and the transposon. The numeric IDs are used to mark distinct read pairs. **e)**, correlations between CiMULE1 (left) and CiMULE2 (right) IL counts and sequencing read abundance in 127 sweet orange accessions. FPKM, fragments per kilobase region per million mapped read pairs. R is the Pearson correlation coefficient and p denotes the p values of the correlation tests. The FPKMs of CiMULE1 or CiMULE2 were calculated using the total read pairs mapped in the shared homologous regions among all its members. The weighted IL counts were calculated by adding the number of the fixed ILs and the non-fixed IL count weighted by the mutant ratios identified in the NGS data. **f)**, IL recall rates of different types of transposon types. Four classes of DNA-type transposons, CACTA, Harbinger, hAT, and MULE, on retrotransposon class LTR, are included in the analysis. Only insertion loci with both upstream and downstream shortest unique K-Mers shorter than 55 bp were subjected to IL recall tests. Mean IL recall rates are labeled using white fonts, and a, b on top of the boxes indicate statistically significantly different groups (FDR<0.01). The center line, median; fork, mean; boxes, first and third quartiles; whiskers, 5th and 95th percentiles.

To address challenges of TE insertion locus (IL) detection brought by the short length of NGS reads and the repetitive nature of transposons (see Methods and Supplementary Note 1), we first used a modified diploid SWO assembly DVS (8) as reference, which allowed distinguishing allele-specific insertions (Figure 1b). Then, we carried out haplotype diversity analyses on the transposon family members (Figure S1 and Supplementary Note 2), based on which we turned repetitive transposons into standalone unique sequences (Figure 1b). This strategy solved the non-unique mapped reads overlapping the TE terminal regions (Supplementary Note 2). Furthermore, we required split-mapped reads for identifying novel ILs, while IL-informative reads (Figure 1c,d), either those split-mapped reads or disconcordantly mapped read pairs, were used for recalling known or identified ILs in accessions, which led to high IL accuracy and high recall rate of known ILs even with low sequencing depth (see Methods and Supplementary Note 1).

We assessed the power of our approach using several realistic datasets. First, the accuracy and recall rates of our NGS pipeline were evaluated using data from three Valencia trees, DVS, T19, and T78, whose Pacbio continuous long reads and NGS data were both available^2^. The CLR and NGS datasets were sequenced using distinct branches in different years, therefore we ignored low (<80%) mutant ratio (mutant-read count / totaling overlapping read count) insertions that were possibly branch-specific in comparison. The results showed that CiMULE1 and CiMULE2 have produced novel ILs among them. For ILs detected using CLR data, 94.8% (55/58 in DVS), 100.0% (19/19 in T19), and 100.0% (19/19 in T78) were recalled on CiMULE1 using our NGS-based approach, and no false positives were detected (Table S4). On CiMULE2, 100.0% recall rates and 100% accuracy were achieved on all three datasets (Table S4). Second, we verified partial of the cultivar group tag-ILs detected in this study, which are only present in all accessions of a specific cultivar group (see following section and Table S5). Analyses showed that all 24 Navel tag-ILs can be found in the Pacbio CLR reads of navel orange var. ‘Gannanzao’^14^ (Table S6). Moreover, we sequenced two Valencia sweet oranges using Pacbio circular consensus sequencing (CCS) and successfully identified all four Valencia tag-ILs (Table S6).

Third, our IL scanning method only utilized dis-concordantly and split-mapped reads overlapping with TE terminals, therefore the read abundance of the TE internal regions could be used to test the reliability of detected IL count. The results (Figure 1e) showed that the number of genomic ILs is significantly correlated with the read abundance for CiMULE1 (Pearson’s r=0.92 and p=2.7E-54) and CiMULE2 (Pearson’s r=0.97 and p=6.0E-78), the two TE families with the largest number of novel ILs. Finally, we tested the presence of ancestral TE insertions in 127 SWOs that were inferred to exist in the most recent common ancestor of analyzed accessions (Supplementary Note 3). As a result, 96.0%, 96.8%, 97.0%, 98.1%, and 98.4% of the ancestral ILs of the CACTA, Harbinger, hAT, LTR, and MULE type transposons were detected in the accessions (Figure 1f). The ancestral IL detection rate of LTR was significantly higher than three (CACTA, Harbinger, and hAT) of the four DNA transposon types while not statistically different (p=0.92) from MULE, suggesting potentially differential transposition mechanisms. Overall, these tests showed that our approach can faithfully unravel genomic TE insertions using NGS data.

### Highly active transposons and discrimination of all SWO cultivars

Then we investigated the transposition activities of the active TE families. We surveyed the ILs of the 34 TE families in 127 SWO, 57 Mandarin, and 39 pummelo accessions (Table S3), among which the SWO accessions were clonal progenies of a single hybrid while all the pummelo and Mandarin accessions had distinct genotypes from hybridization. As a result, the SWO, mandarin, and pummelo accessions had significantly different lengths of evolutionary divergence. Our method detected 4,850 SWO-specific ILs, 12,309 mandarin-specific ILs, and 4,900 pummelo-specific ILs in them (Figure 2a and Table S7).

**Figure 2.**
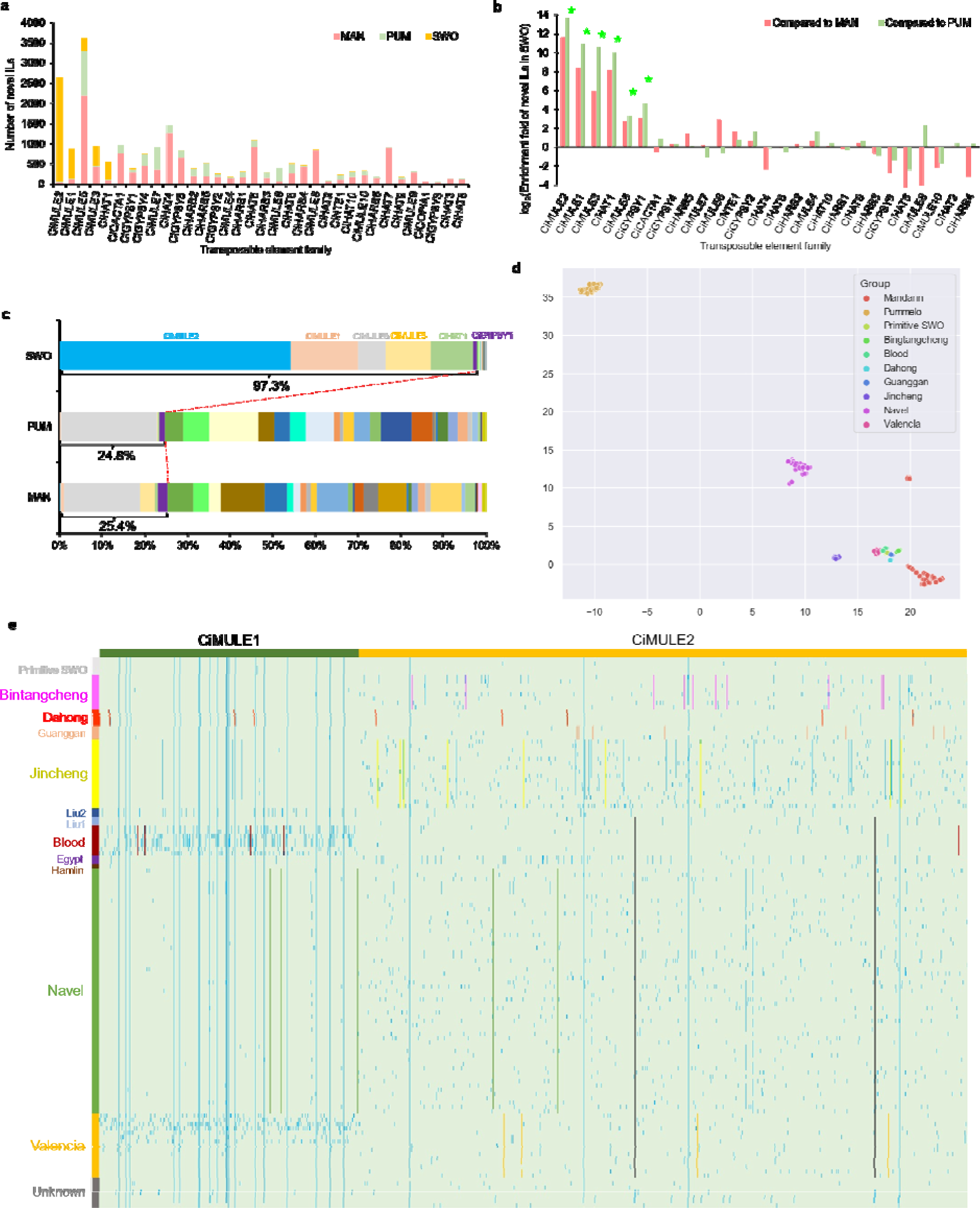
Diversity of transposon insertions in sweet orange genomes. **a**, stacked bar plots of novel IL counts from the 34 TE families in sweet orange (SWO), pummelo (PUM) and mandarin (MAN). **b**, enrichment of novel ILs in sweet orange. Green stars denote novel ILs being significantly (FDR < 0.05 b Chi-square tests) overrepresented or underrepresented in sweet orange compared to both mandarin and pummelo. The test was carried out against the median ratios of the 34 transposon families among the three species. **c**, proportions of novel IL contribution of the 34 TE families in the three species. Red dashed lines and the percentages mark the overall novel IL contributions of the six boosted TE families. **d**, t-SNE (t-distributed Stochastic Neighbor Embedding) projection of SWO cultivars, pummelo and mandarin accessions’ IL-based genotypes to lower dimensions. **e**, CiMULE1 and CiMULE2-based accession IL fingerprints and cultivar group tag-ILs (colored lines). Each light green or blue bar denotes the absence and presence of the IL (line) in a corresponding accession genome (row). For each cultivar group on the left, its tag-ILs were colored the same as the cultivar group. The two non-China origin ILs were colored black.

To test if TE families had significantly different transposition activity levels among the three species, we carried out IL enrichment tests for the TE families among SWO, mandarin, and pummelo. The test was carried out against the null hypothesis that the majority (median) of TE families remained similarly active among the three species. This analysis unraveled six boosted TE families with significantly enhanced transposition activities in SWO compared to those in mandarin and pummelo (Figure 2b and Table S7), including 5 DNA transposon families (CiMULE1,2,3,5 and CihAT1) and 1 LTR retrotransposon family (CiGYPSY1). These six TE families have contributed 97.3% of novel ILs to SWO, but only 25.4% of mandarin-specific ILs and 24.8% of pummelo-specific ILs (Figure 2c). CiMULE2 alone has produced a majority (53.5%, 2,596) of the novel ILs in SWO, followed by CiMULE1 (751), CiMULE3 (512), CihAT1 (468), CiMULE5 (338), and CiGYPSY1 (52). Rough estimation showed that CiMULE2 has been ∼3,095.5- and ∼8,974.2-times active in SWO compared to in Mandarin and pummelo, respectively. SWO also had 7.4-to 1,397.9-fold novel IL enrichment on the rest five boosted TE families.

The high diversity of ILs (38.2 novel IL per accession genome) allowed for solving the difficult puzzle of molecular discrimination of all SWO cultivars. All analyzed SWO cultivar groups were separated via T-SNE (t-distributed Stochastic Neighbor Embedding) projection of their IL-based genotypes (Figure 2d). Cultivar group tag-ILs can be used to distinguish different cultivar groups and are potentially involved in phenotype formation of cultivars but have seldom been identified in SWO before^5, 15^. Here, we have found abundant tag-ILs for all analyzed SWO cultivar groups, including Navel, Valencia, Blood, Egypt, Jincheng, Dahong, Guanggan, and Bingtangcheng (Tables S4 and S8). The most active CiMULE2 has contributed 1 (Blood) to 11 (Bingtangcheng) tag-ILs to all eight analyzed cultivar groups. Though LTR type CiGYPSY1 has produced the fewest number of novel ILs, it has yielded tag-ILs for 7 cultivar groups. CiMULE1, CiMULE3, CiMULE5, and CihAT1 have also contributed tag-ILs to 4, 1, 3, and 4 cultivar groups, respectively. Moreover, 126 of the 127 analyzed sweet orange accessions can be discriminated against using accession-specific ILs from all TE families or even CiMULE2 alone except for two Xuegan cultivars (Figure 2e and Table S9). The rest five boosted TE families have also produced tag-ILs that could identify specific SWO cultivars (Table S10). Our IL scanning approach also enables known IL recalling with statistical confidence by using ILs in non-repetitive regions (see Methods). Overall, our high-accuracy IL scanning approach and the six boosted TE families have allowed fingerprinting of SWO cultivars.

### Sweet orange breeding and dispersion history told by transposons

To learn the history of SWO, researchers previously constructed an unrooted phylogenetic tree of asexually propagated SWO accessions using somatic single nucleotide mutations^4^. However, it failed to reconstruct the early diverging and dispersion events and did not have robust support for major cultivar groups such as Navel or Valencia oranges. Here, we successfully used the identified TE ILs to reconstruct the lineage development and dispersion history of SWO. As shown in Figure 3a, our IL-based rooted phylogenetic tree has provided robust support for major SWO (sub)cultivar groups including Navel, Valencia, Blood, Bingtangcheng, Jincheng, et al.. And it supports three major lineage-dispersion events in SWO breeding history (Figure 3b). The first dispersion event gave rise to primitive SWOs, including four accessions that were ancestors of UKXC, Taoyecheng (TYC), Xuegan, and all the rest analyzed cultivars (referred to as modern SWOs), respectively. UKXC, an accession from Hunan province, is the closest to the inferred ancestor of SWO, differing by only one CiGYPSY1 IL. TYC originated in Hubei and Xuegan originated in Guangdong, which are neighboring provinces of Hunan.

**Figure 3.**
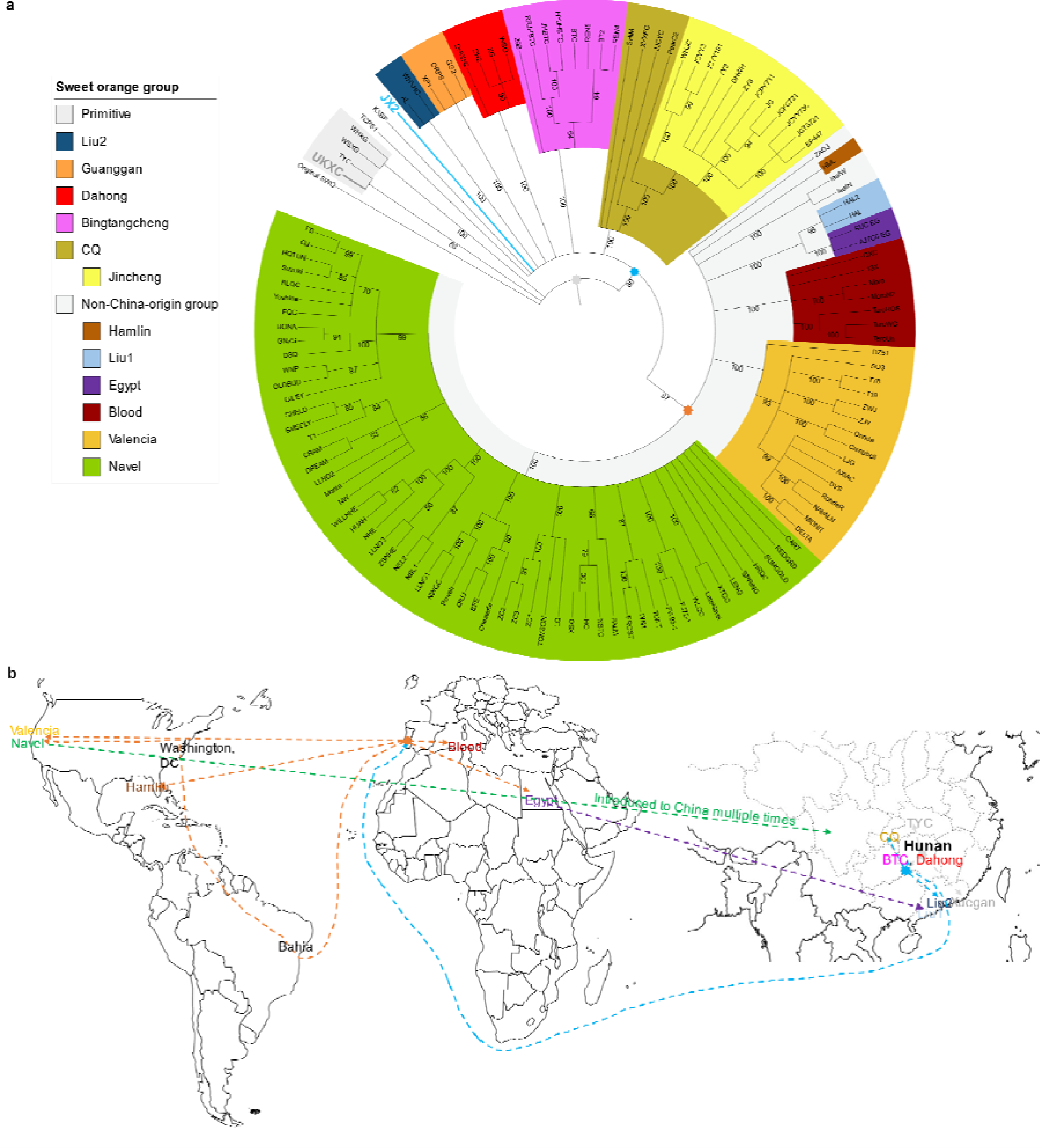
Sweet orange lineage development and dispersion history. **a)**, Transposon insertion loci-based sweet orange breeding history. The phylogenetic tree was constructed using 1,150 ILs from the active transposon families detected in at least two accessions. The root (original SWO) of the tree was a hypothesized accession with none of the SWO-specific ILs. In the cladogram, values on the branches are the posterior probabilities (%) from Bayesian inference, and we have deleted branches with <50% posterior probabilities. Distinct background colors indicate different cultivar groups as denoted in the legend. The silver, light blue, and orange polygons represent the inferred ancestral lineages distributed in the three major dispersion events, respectively. **b),** The dispersion of sweet orange. To better display the dispersion of sweet orange in China, the partial world map has an enlarged southern China region. Straight dashed lines point from the ancestral lineage location to the offspring location with unknown transmission route information. Based on historical documents, curved dashed lines show the rough transmission route prior to the origin of a cultivar group.

The second major dispersion, estimated to have occurred between 1470 and 1533 AD (see Methods), distributed the common ancestor of modern SWO in China. This inferred ancestor carried four ILs (referred to as modern-ILs) from CiMULE2, CiMULE5, CiHAT1, and CiGYPSY1, respectively, which exist in the genomes of modern SWOs but not in any primitive SWO. This dispersion event was rapid and led to China-local cultivar groups, including Chongqing (CQ), Dahong, Guanggan, and Bingtangcheng, as well as the common ancestor of non-China origin groups, which commonly shared the four modern-ILs but not any other novel ILs with each other group. JX2, a local cultivar from Jing Xian in Hunan province, is the closest to this ancestor, differing by only four novel CiMULE5 ILs. Therefore, Hunan province, which is also the origin location of Bingtangcheng and Dahong and adjacent to Chongqing, can be inferred as the cradle of SWO and the center of its first and second major dispersion events (Figure 3a).

The third dispersion event led to all SWO groups that developed out of China (referred to as non-China groups), including Navel, Valencia, Blood, Hamlin, and Egypt accessions. Consistent with the single nucleotide mutation-based phylogeny^16^, the TE-based phylogenetic tree also supports the monophyly of non-China origin cultivars (Figure 3a). The distributed ancestral lineage should harbor two novel CiMULE2 ILs (called NC-ILs) shared only by all accessions from non-China groups. According to the documentary, this dispersion event most likely occurred soon after the first modern sweet orange (Portugal orange) was introduced to Portugal around 1635^17^. The non-China groups then developed independently after the transmission since different groups do not share any other novel IL. The phylogenetic tree also supports that a few non-China origin lineages were introduced back to China, including several Navel orange lineages and the ancestor of two accessions (Liu1) misrecognized as traditional Liucheng (Figure 3b). Liu1 harbors the two NC-ILs and shares several ILs specifically with lineages from Egypt, implying a Mediterranean area origin. Two other accessions (Liu2) named Liucheng do not have the NC-ILs and should be authentic local accessions from Guangdong.

### Dynamics of transposition activity in sweet orange

Based on the TE ILs and cultivar breeding history, we unraveled the transposition activity levels (estimated using IL accumulation rate) of these six boosted TE for different phylogenetic paths in Figure 3a. The transposition activity levels of the original SWO plant were then inferred based on the primitive SWO accessions. Four (excluding CiMULE5) of the five boosted DNA TE families had significantly lower transposition activity levels in primitive SWOs compared to modern SWOs (Figures 4a and S2, and Table S11). We observed zero IL accumulation rate of the five DNA TE families in at least half of the primitive SWO accessions. UKXC, the SWO accession closest to the original SWO in respect of ILs, has not accumulated any ILs of the five boosted TE families. Meanwhile, CiGYPSY1 had novel IL accumulation in all primitive SWOs. Overall, the six boosted TE families were inferred with low (<0.01 IL/year) or zero transposition activity levels in the original SWO plant.

**Figure 4.**
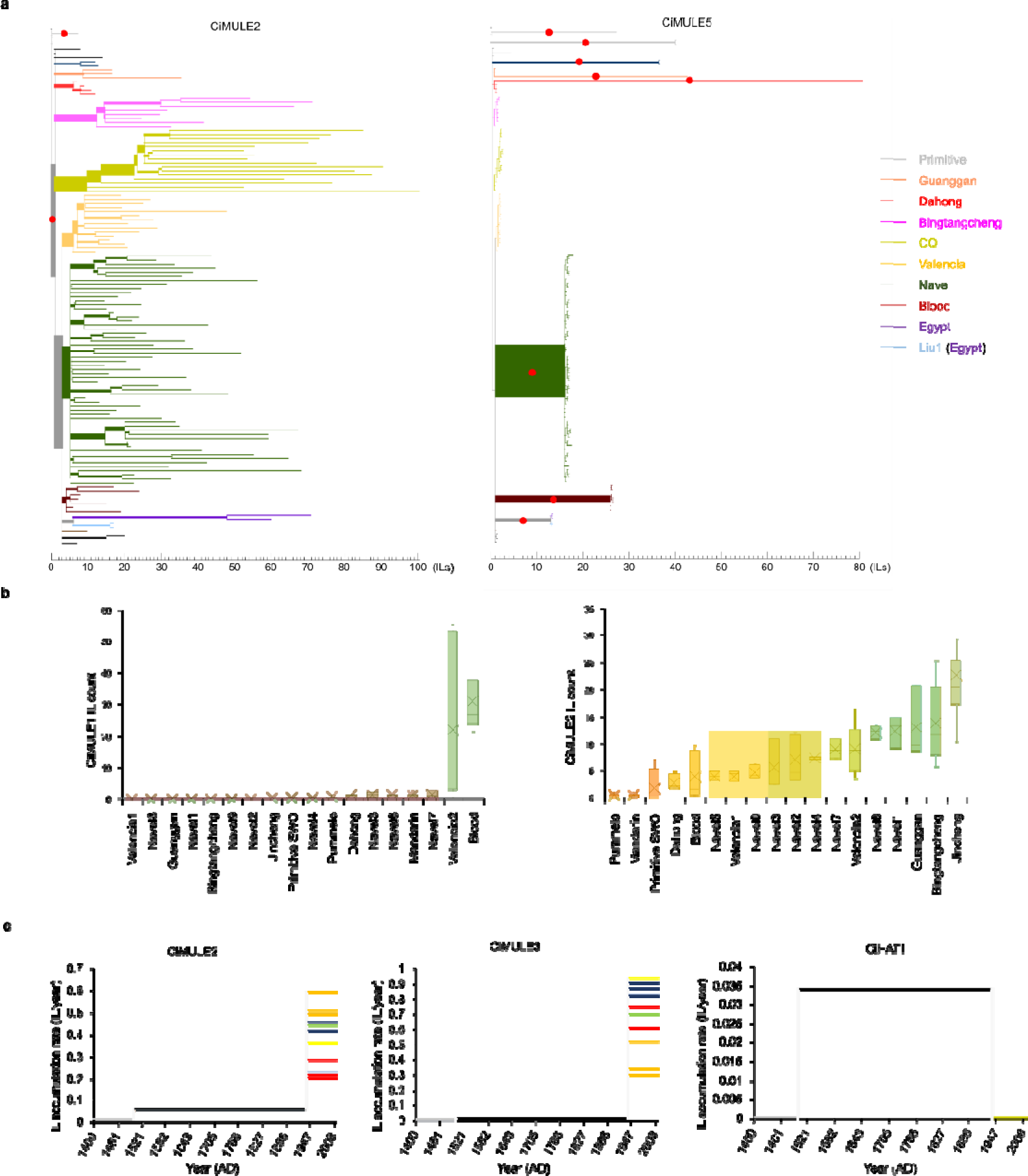
Transposon insertion accumulation in sweet orange lineages. **a)**, Insertion locus (IL) accumulation in phylogenetic paths. The tree topology is the same as Fig. 1a, and the branch lengths are proportional to the numbers of accumulated novel ILs. In the scale bars, each minor mark represents one novel IL, and branches shorter than one minor scale are only used to display tree topology. Branches with inferred transposon activation events are marked with solid red circles. Same as Fig. 1a, the line colors denote distinct cultivar groups. **b**, Distribution of novel ILs (CiMULE1 and CiMULE2) in mandarin, pummelo, and distinct sweet orange phylogenetic clades (Table S16). The IL counts were weighted b mutant ratios and the reciprocal of the number of accessions sharing the ILs. Each background color indicates a statistically (by one-way ANOVA test) homogenous subset. In the boxplots: the center line, median; fork, mean; boxes, first and third quartiles; whiskers, 5th and 95th percentiles. **c)**, Mean IL accumulation rates in Jincheng accessions and their common ancestor. For CiMULE2 and CiMULE3, differentially colored lines indicate different phylogenetic sub-clades in Jincheng. CiHAT1 has been silent in all Jincheng accessions which have been merged into a yellow line.

Each of the five boosted DNA TE families has been activated (arising to accumulating above 0.01 ILs/year) independently in the SWO breeding history (Figures 4a and S2). The activation of CiMULE2 has occurred twice, in the common ancestor of modern SWOs and TYC ancestors. Activation of the rest four boosted DNA TE families has each been observed on at least one phylogenetic path leading to a major cultivar group. CiMULE1 and CiHAT1 were independently activated in the ancestors of Dahong, Jincheng, Navel, and Blood; CiMULE5 activation occurred in the ancestors of Navel, Blood, and Egypt groups, respectively; CiMULE3 was activated in the ancestor of the CQ group. We also observed their independent activations in individual lineages (Table S11). For instance, CiMULE1, CiMULE3, and CihAT1 were not active in the common ancestor of Valencia, but it has become active in several independently selected Valencia lineages (Figures 4a and S2). Though CiGYPSY1 has maintained similar IL accumulation rates in most SWO lineages, it also exhibits higher transposition activity levels in the phylogenetic paths leading to the Bingtangcheng ancestor and several other accessions (Figure S2 and Table S11).

After TE activation, we observed a significant increase in IL accumulation rates within decades or centuries in different SWO lineages. In modern SWOs, the summed IL accumulation rates of the five boosted DNA TE families varied from 0.010 ILs/year (JX2) to 1.385 ILs/year (JCTS721) (Table S11). We have observed up to 20-fold difference in CiMULE2 IL accumulation rate among modern SWO clades (Figure 4b). In Guanggan, Bingtangcheng, Jincheng, Valencia, Navel, and the Egypt group, independently selected lineages had significantly enhanced CiMULE2 transposition activities compared to their respective group ancestors. Furthermore, the results showed that sub-clades of a cultivar group could obtain significantly increased IL accumulation rates within down to tens of years. For instance, the IL accumulation rates of CiMULE2 and CiMULE3 have increased up to 3-fold in the sub-clades of the Jincheng group (Figure 4c), which has a documented history of ∼80 years. Navel orange sub-clades had up to 5-fold CiMULE2 IL accumulation rate increase in the past ∼147 years (Table S11). Similar cases also happened in the Blood group on CiMULE1 and CiHAT1 and in the Bingtangcheng, Blood, Valencia, and Egypt groups on CiMULE2.

Transposon silencing (dropping to <0.01 ILs/year accumulation rate), which is important for genome protection in eukaryotes ^18, 19^, has functioned differently on the five boosted DNA TE families. We have observed a <1% silencing ratio of CiMULE2 (only silenced in JX2) in modern SWO accessions (Figure 4a and Table S11). CiMULE3 has a zero percent silencing ratio in Jincheng (Figure S2). In contrast, CiMULE5 became silenced in almost all cultivar groups (Navel, Blood, and Egypt) after its activation in the groups’ ancestors (Figure 4a). CiMULE1 and CiHAT1 have been silenced in no less than 50% of Jincheng, Dahong, and Navel oranges, but none of the Blood oranges (Table S11).

Transposition activity could be related to transposase expression, which is the prerequisite of transposition for DNA type TEs^20^. CiMULE2 is the only autonomous TE family among the six boosted TE families. The expression of CiMULE2 transposase in 629 SWO, 39 mandarin, and 171 pummelo transcriptomes were analyzed to learn its relationship with transposition activity (Table S12). Significantly higher expression levels of CiMULE2 were detected in most SWO cultivars/cultivar groups compared to those in Xuegan (primitive SWO group), mandarin, and pummelo (Figure S3a). The expression of CiMULE2 is diversified among the SWO groups corresponding to its transposition activity levels. We have observed significant positive correlations between CiMULE2 expression abundance and the IL counts in the cultivar groups (Figure S3a, b). Transcriptome analysis also showed that CiMULE2 utilizes many external transcription starting sites (Tables S13 and S14). As a result, every new IL of CiMULE2 has the potential to increase its expression and transposition activity, which explains the highly diversified CiMULE2 transposition activity levels observed among SWO cultivars.

### Contribution of transposons to sweet orange cultivar formation

The boosted TE families have contributed to the genetic diversity of SWO cultivar groups (Figure 5a). We found a total of 2,775 ILs including 2,424 from the boosted TE families located in genic or promoter regions of 3,185 genes in the 127 SWO accessions (Table S15). These genes are significantly overrepresented in plant development processes and phytohormone signaling pathways (Figure 5b), which are closely related to citrus horticultural traits. The impacted genes in different sweet orange cultivar groups are unevenly distributed in the functional categories (Figure 5c). Moreover, independent ILs impacted 355 genes twice or more times in SWO accessions (Figure 5d), which is enriched by 5.1E4-fold (*p*<1E-99) compared to that by random and stands as a strong signal of selection. And these IL-impacted genes are valuable candidate horticulture traits-related research targets.

**Figure 5.**
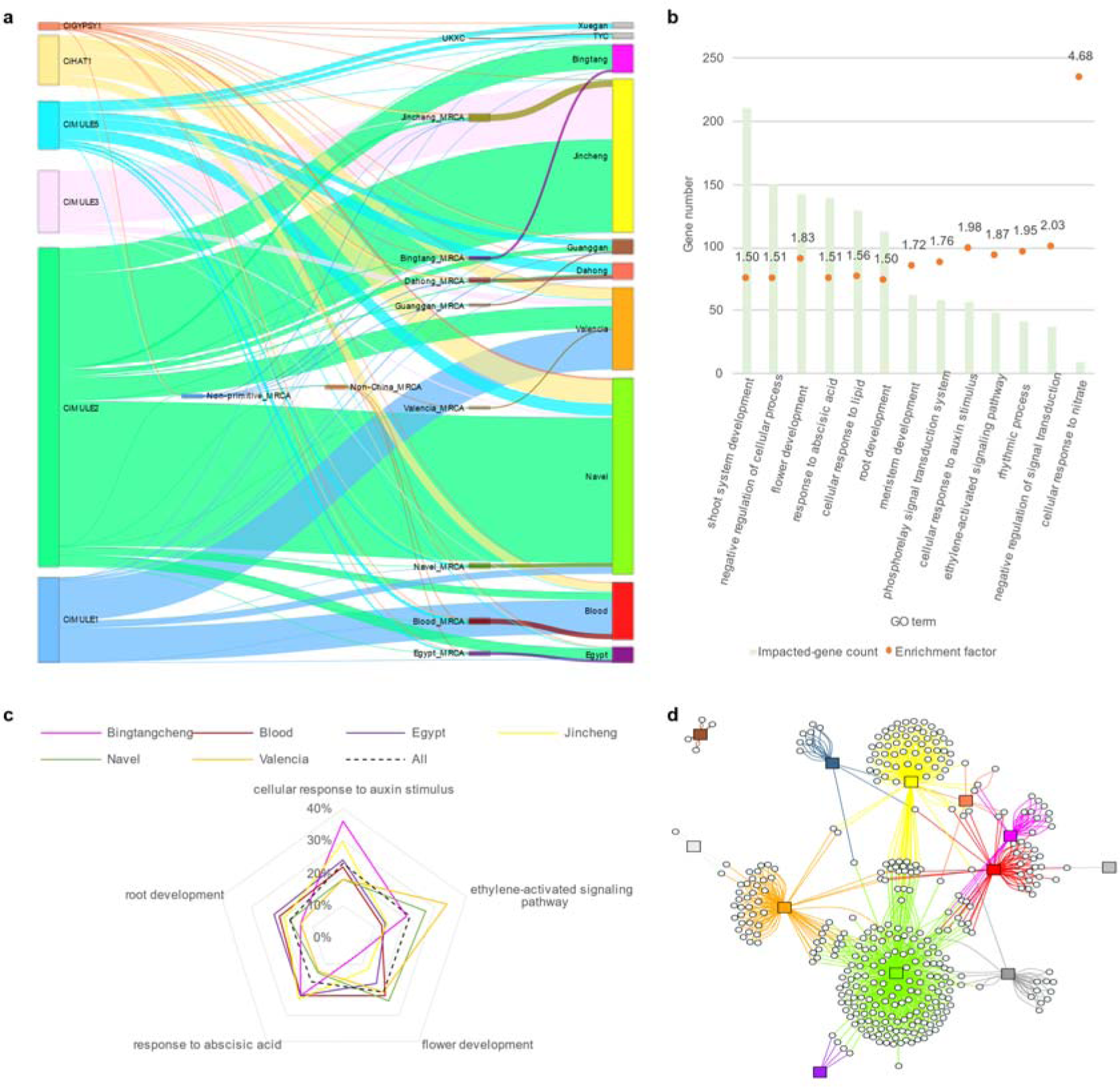
IL contribution of the boosted TE families to sweet orange groups and impacted genes/pathways. **a)**, Novel insertions from the six boosted TE families in sweet orange cultivars. The width of the links is in proportion to the TE insertion counts. The total collateral links in this graph represent 4,650 TE insertions. The link colors follow the source TE families or the most recent common ancestors (MRCA) from which the insertions have been inherited. **b)**, Gene ontologies (GOs, biological processes only) enriched in TE insertion-impacted genes. **c)**, Relative abundance of TE insertion-impacted genes in five horticultural traits related to GO terms in sweet orange groups. The impacted gene count of each GO term was normalized by its total gene count in the reference genome, and then the relative abundance of impacted genes was calculated for each GO term. All denote all the 127 analyzed sweet orange accessions. **d)**, Distribution of genes impacted by at least two independent TE insertions events in sweet orange. Each circular node denotes a gene locus and each rectangular node represents a sweet orange cultivar group (colored the same as in panel A). Three hundred and thirty-five genes impacted more than one independent TE insertions are drawn in this graph. Each link (colored the same as the connected sweet orange group) denotes the gene (either allele) being impacted by one TE insertion in the connected cultivar group.

The SWO group tag-ILs have potentially affected the function of 138 genes (Table S15). Two ILs impacting both alleles of one gene in the same lineage are referred to as biallelic ILs. We found that four SWO cultivar groups owned 7 pairs of biallelic tag-ILs (Table 1). Two Jincheng tag-ILs are located in the 5’-UTR and upstream region of *CsDLO* alleles, which are orthologous to *AtDLO1*, *AtDLO2*, and *AtDMR6* that act as immunity suppressors in *Arabidopsis*^21^, are potentially related to the good post-harvest storage stability of Jincheng. We also detected biallelic ILs in individual sweet orange cultivars. Moro blood orange has biallelic CiMULE2 ILs in intron and upstream regions (within 2 kb if not specifically described) of *CsBRC2* alleles. *AtBRC2* and *AtBRC1* prevent axillary bud outgrowth in *Arabidopsis*^22^, and disruption of *BRC1* ortholog in citrus converted thorns into branches^23^. These *CsBRC2*-impacting biallelic ILs putatively contribute to the branchy characteristics of the Moro accession. In Campbell Valencia orange, which is more vigorous, thornier, larger, broader-topped, and slower to come into bearing than ordinary Valencia orange is^3^, two CiMULE1 ILs are detected in the coding regions of two alleles of *CsCYP90C1*, which is involved in brassinosteroid biosynthesis^24^.

**Table 1.** Representative group tag transposon insertions.

| Transposon insertion(s) | Biallelic mutations | TE family | Group tag-IL | Impacted gene |
| --- | --- | --- | --- | --- |
| chr3A:5225560-5225561 / chr3B:5278735-5278736 | Yes | CiGYPSY1 | Bingtangcheng | AT5G24080 ortholog |
| chr8A:25898219-25898229 / chr8B:27879542-27879546 | Yes | CiMULE2 | Bingtangcheng | *CsALPHA-ADR* |
| chr3B:14705892-14705903 | No | CiMULE2 | Bingtangcheng | *CsNCED1* |
| chr2A:32769747-32769757 | No | CiMULE2 | Bingtangcheng | *CsNHX6* |
| chr6B:23447469-23447479 | No | CiHAT1 | Blood | *CsABAH1* |
| chr6A:16294504-16294508 | No | CiGYPSY1 | Blood | *CsAUX1* |
| chr4A:28130579-28130620<br>/ chr4A:28130730-28130736 | Yes | CiMULE5 | Blood | *CsAPC2* |
| chr1B:26750047-26750053 | No | CiMULE5 | Blood | *CsCML7* |
| chr3B:4333882-4333897 | No | CiMULE2 | CQ | *CsCML37* |
| chr7B:11903747-11903749<br>/ chr7A:11877823-11877828 | Yes | CiMULE2 | CQ | *CsDLO* |
| chr1A:21260169-21260188 | No | CiMULE2 | CQ/Jincheng | *CsFRL4A* |
| chr4A:24354360-24354361<br>/ chr4B:23240217-23240223 | Yes | CiMULE2 | CQ/Jincheng | AT2G03780 ortholog |
| chr4B:25528500-25528511 | No | CiMULE2 | CQ/Jincheng | *CsPGL3* |
| chr8A:23003963-23003973<br>/ chr8B:25316854-25316868 | Yes | CiHAT1 | CQ/Jincheng | *CsYAB2* |
| chr9A:31585671-31585680 | No | CiHAT1 | Dahong | *CsPIRL6* |
| chr1A:4433776-4433788 | No | CiMULE2 | Dahong | *CsYAB5* |
| chr4A:21899887-21899888<br>/ chr4B:21134956-21134957 | Yes | CiMULE2 | Guanggan | AT4G19060 ortholog |
| chr9A:11153462-11153473 | No | CiMULE2 | Guanggan | *CsCML36* |
| chr5A:35277942-35282999 | No | CiMULE2 | Valencia | *CsSERK2* |
| chr8A:28575073-28581996 | No | CiMULE2 | Valencia | AT2G28710 ortholog |
| chr3A:1797300-1797310 | No | CiHAT1 | Navel | *CsMURE* |
| chr1B:2608945-2613410 | No | CiHAT1 | Non-primitive SWO | AT2G25670 ortholog |
| chr4A:21347337-21347342 | No | CiGYPSY1 | Egypt | *CsSIGA* |
| chr1A:7072375-7072385 | No | CiMULE2 | Egypt | *CsPAP2* |
| chr4A:15316629-15316635 | No | CiMULE5 | Egypt | *CsBAM2* |
| chr3A:41996808-41996809 | No | CiMULE5 | Egypt | *CsCLASP* |
| chr2A:26415866-26415877 | No | CiMULE1 | Blood | *CsUFGT* |
| chr5A:26649742-26654799 | No | CiMULE2 | Valencia | *CsGID1C* |

Non-biallelic ILs could also be functionally important in sweet orange cultivar group formation (Table 1 and Table S15). In blood orange, a tag CiMULE1 IL is in the 5’-UTR region of the *CsUFGT*, which plays an important role in the biosynthesis of anthocyanins and is related to flower and fruit colors in plants ^25, 26^. Our analyses of public RNA-Seq data showed that this IL has significantly increased the allelic expression of the pummelo-origin *CsUFGT*s in blood oranges compared to other SWO cultivars, indicating it has contributed to the anthocyanin accumulation together with another retrotransposon insertion in the promoter region of the *Ruby* gene ^5^. One Bingtangcheng tag IL is in the 5’-UTR of *CsNCED1*, a key gene in ABA biosynthesis and related to fruit coloring and ripening ^27^. Another Bingtangcheng tag is IL in the 5’-UTR of *CsNHX*, which has been related to the low acid characteristics of Bingtangcheng ^4^. Two Valencia tag-ILs are located upstream of *CsGID1* and *CsSERK1*, which play important roles in gibberellin ^28^ and brassinosteroid ^29^ signaling, respectively. Several other cultivar group tag-ILs are located upstream of or inside genes related to phytohormones, including brassinosteroids, gibberellins, abscisic acid, and auxin (see Supplementary Note 4).

## Discussion

Transposable elements (TEs) have traditionally been regarded as "selfish genes" or "junk DNAs" considering their movements are mostly neutral or detrimental to organisms^30^. In evolution, selective pressures continuously shape the activity levels of transposons, with negative selection generally favoring individuals with efficient silencing mechanisms such as RNA interference, DNA methylation, and histone modifications^31, 32^. Nevertheless, recent studies unraveled that transposons have played crucial roles in shaping genomes in evolution^33, 34^, as well as producing genetic variations that contribute to phenotypic diversity of domesticated species^35^ such as maize^36^, citrus^37, 38^, and grape^39^. However, TE activities are hard to predict or interpret since they are influenced by both intrinsic and extrinsic factors of organisms with high complexity. Diversified types of TE families^40^, either belonging to retrotransposons (type LJ) or DNA transposons (type LJ), have varied transposition mechanisms^41^ and exhibit distinct transposition activities. As a result, TE activity dynamics and the full scenery of their insertions in most horticultural species remain poorly understood.

Our research reveals that TE insertions are present as tag mutations for all SWO cultivar groups. Moreover, they can distinguish over 99% of SWO accessions and are significantly enriched in affecting development and phytohormone signaling pathways. Our results show that the high bud sport selection frequency of SWO could be explained by the activation of boosted TE families in SWO. The overall IL accumulation rates in modern SWO are significantly higher than those in the original SWO. This high transposition activity level has accelerated SWO breeding and played an essential role in the modern SWO industry. Most significantly, the activation of CiMULE2 most likely has been the prerequisite for the development of many modern SWO cultivar groups.

Our results imply that transposons could have served as important mutation rate adjustors in evolution. Species explosions are usually accompanied by transposon family expansion after dramatic environmental changes, which result in high positive selection pressures^42^. It has long been hypothesized that positive selections are prone to select lineages with higher mutation rates indirectly in asexual organisms, which has only been proven in microorganisms^43, 44^. In breeding by selection of somatic mutants, a minimum mutation rate is necessary for a genotype to accumulate enough mutations within a certain time (Figure S4). SWO has been the subject of bud sport and nucellar seedling selection for over 500 years^3^, resembling continuous adaptive evolution in nature^45^. The increased transposition activities in SWO lineages are inferred to be the byproducts of repeated artificial selections of desired traits^12^. The dramatic increase of the IL accumulation rates in SWO has occurred in a short time (tens to hundreds of years), suggesting the involved TE families are very sensitive to the selections and have functioned as mutation accelerators under human selection. Compared to other mechanisms altering mutation rates such as alterations in DNA replication and repair mechanisms, TEs are much more controllable in promoting mutation rates and less harmful in the long term, since many organisms have developed epigenetic mechanisms to silence the transposons^46, 47^.

In conclusion, transposons, especially the six boosted TE families, have driven the selection and diversification of modern sweet orange cultivars. Sexual propagation and continuous bud sport selections can lead to lineages with higher transposition activity levels in a short time, which could have been utilized without being noticed in many horticultural species.

## Materials and Methods

### Searching for active transposons among SWO assemblies

To identify active transposable elements (TEs) in SWO, we obtained genome assemblies of 11 SWOs^2, 4^ (Table S1) from the NCBI assembly database. we used DVS^2^ as the reference genome and aligned the rest ten assemblies to it using Minimap2 v2.17^48^. Large (>100 bp and < 20 kb) indels were called using the paftools.js script supplied with Minimap2 v2.17, and the inserted or deleted sequences were output in FASTA format. We screened the indel sequences homologous to TEs annotated in the DVS genomic annotation with blastn E-value <1E-3. The screened sequences were clustered using CD-HIT v4.8.1^49^ with ≥90% nucleotide similarity, and TE family members were required to have intact 100 bp terminals (≥90% similarity and <10bp terminal truncations) at both ends. The TE family members were further searched in the 11 assemblies by blastn using the same criteria. As a result, we found 34 TE families to be active with no less than two distinct insertions among the 11 assemblies. Then the TE families were classified into different retrotransposon or DNA transposon types using DeepTE^50^. The detected members of TE families were further clustered into subgroups using CD-HIT v4.8.1 with no less than 99% nucleotide similarity and 99% minimal alignment coverage (Figure S1). ClustalX v2.1^51^ was applied in the multiple sequence alignment. The haplotype diversity in the 50 bp aligned regions at both terminals of TE families was summarized from the multiple sequence alignments.

### Analysis of factors impacting TE insertion scanning

We delved into accurate TE IL scanning for SWO using whole-genome NGS data. Sequencing reads/read pairs mapped across IL boundaries are used for precisely locating and recalling the ILs (Figure 1c,d). Multiple alignments (primary alignment MAPQ=0) indicate ambiguous alignment targets, therefore we only used unique read alignments for IL detection. To assess the impact of the factors on IL detection, we tested different combinations of factors described in Figure 1c,d using in silico simulations. The results (Figure S5 and Supplementary Note 1) showed that the capability to recall (know the existence) and precisely locate an IL using NGS data are affected by sequencing depth, mutant ratio of the IL, the shortest unique K-Mer lengths (SULs) at the ends of the scanned TEs (SUL2 and SUL2’ in Figure 1c,d) and surrounding the ILs (SUL1 and SUL1’), the sequencing library insert size, and the sequencing read length. For either simulation or realistic data, at least two non-duplicate read pairs including at least one split-mapped read were required for identifying novel ILs, while ≥2 IL-informative reads pairs of either type would recall an IL in an accession (Figure 1c,d).

### SWO transposon insertion scanning

Accordingly, optimizations were made and the procedures of the IL scanning mainly included (detailed in Supplementary Note 1): (1) identification of >100 bp continuous regions in the pseudo-haplotype chromosome set DVS_B of DVS with identical allelic regions in DVS_A, then masking these regions using the ambiguous base N except for their terminal 50 bp regions; (2) masking all members of analyzed TE families in DVS, and adding standalone copies of TE cluster representatives with duplicate terminal 50 bp regions masked except for one (Figure 1b); (3) analyzing the length distributions of the unique k-mers in the masked reference DVS genome, the standalone TE member terminals, and the surrounding sequences of the ILs detected from the 11 SWO assemblies (Figure 1a); (4) masking 10 bp regions surrounding the allelic regions of reference ILs to avoid false positive calls; (5) mapping the NGS data to the modified DVS reference genome using bwa-mem; (6) detecting two types of IL supporting the read pairs (only unique alignments are taken into count): A) read pairs including a read split-aligned to, and B) read pairs without split-mapped reads but dis-concordantly mapped to both the standalone transposon sequences and the chromosomes (Figure 1d); (7) ILs with both type A and type B reads in at least one sample were regarded as high-quality ILs, which were subjected to genotyping in all analyzed accessions. The precise coordinates of the ILs were inferred using the split-mapped reads; (8) for a type B read pair, the possible range of its IL is inferred based on the read pair orientation and the library insert size, and this range is applied to assign the read pair to a high-quality IL according to their overlapping relationship. When the inferred IL range of a type B read pair is overlapped with two high-quality ILs, it would be assigned to the one with split-mapped reads in the sample or not assigned if no split-mapped read of either IL was present; (9) at least two non-duplicate supporting read pairs (either type A or B) are required to recall an IL in a sample; (10) The number of read pairs from the mutant-type (*C_m_* in Figure 1d) and the wild-type (*C_w_*) cells on each IL are counted in the mutant samples. Then the mutant ratio is calculated depending on whether and how the allelic region (including the 1 kb surrounding region) of the IL is masked: a) when the allelic region is not masked, the mutant ratio: (*C_m_*/2) / (*C_m_*/2 + *C_w_*); b) when the allelic region is fully masked: *C_m_* / (*C_m_*/2 + *C_w_*); c) when the allelic region was partially masked, the read pairs mapped overlapping the allelic region will be re-mapped to DVS_A, and the count of the effective wild-type read pairs (Figure 1d) will be added to *C_w_*, then the mutant ratio is calculated as *C_m_* / (*C_m_*/2 + *C_w_*). In the scripts, BEDTools v2.29.2^52^ was applied in genome masking and genomic interval overlapping analysis, BWA v0.7.17^53^ was applied in read mapping, and Samtools v1.10^54^ was used to read the alignment files.

### Assessments of the IL scanning approach

The Pacbio CLR reads of navel orange var. ‘Gannanzao’ (SRR25380064)^14^ were downloaded from NCBI. Leaves of two Valencia orange trees were collected from the resource nursery of the U.S. Horticultural Research Laboratory in Fort Pierce, Florida, and subjected to DNA extraction and Pacbio CCS (∼30×) sequencing by BGI Genomics, Shenzhen, China. To verify the detected ILs using Pacbio CLR or CCS reads, we used Minimap2 v2.17 to align them to the same modified DVS reference genome applied in NGS-based IL scanning. Uniquely split-mapped reads (MAPQ>=10) overlapping the IL upstream or downstream region and the corresponding TEs were identified as IL supporting reads. At least two such reads were required to verify an IL in an accession.

For each TE family, the normalized sequencing read abundance (FPKM, fragments per kilobase region per million mapped read pairs) was calculated based on the reads mapped to the homologous regions shared by all its members. We then carried out Pearson’s correlation test between the total IL counts (fixed IL counts plus mutant ratio weighted non-fixed IL counts) and the FPKM values, respectively. In the last test, the ancestral ILs were inferred based on their universal presence in the three primitive SWO lineages (Xuegan, UKXC, and TYC).

### IL-based phylogenetic analysis

We classified the ILs as fixed (≥90% mutant ratios) and non-fixed (<90% mutant ratio) according to their mutant ratios in the sequencing data. Only fixed ILs were used in phylogenetic analysis. The genotypes of the analyzed accessions on each IL were detected as 1 (present in accession) and 0 (absent in accession). For SWO cultivar groups including 3 or more accessions, we looked for the tag-ILs present in >90% of the cultivar group accessions and absent in all accessions from other cultivar groups. Only ILs present in at least two SWO accessions were used in phylogenetic analysis. We carried out phylogenetic inference on 127 SWO accessions and two pummelo accessions as outgroups using the Bayesian inference method by MrBayes v3.2.7, with the model assuming equal mutation rates across all ILs, 110,000 generations of MCMC sampling, 1/100 tree sampling frequency and 100 burn-in trees. Branches with less than 50% posterior probabilities were deleted in the phylogenetic tree.

### IL accumulation analysis

The ILs count accumulated in each accession was calculated by summing the total IL counts weighted by mutant ratios. For instance, an IL with mutant ratio (*R_M_*) is only counted as (1 × *R_M_*). We carried out one-way ANOVA (analysis of variance) on IL counts of the five boosted DNA TE families among pummelo, mandarin, and SWO groups/clades. The analyzed SWO groups/clades include the primitive SWO group, the four earliest diverging Valencia accessions (VAL1), and SWO phylogenetic clades including ≥3 accessions and with ≥ 80% posterior probabilities (Table S16). The distributions of the weighted IL counts in pummelo and Mandarin (groups with the largest sample sizes) are right-skewed, thus we performed log_2_ transformations of the IL counts before one-way ANOVA. We then applied the Student-Newman-Keuls post-hoc tests to detect different homogeneous subsets which include groups with non-significantly different means at the α = 0.05 level implemented in IBM^®^ SPSS^®^ Statistics v26 (IBM, Armonk, USA).

The IL accumulation rate on each phylogenetic branch (Table S11) was calculated by dividing the corresponding IL count by the length (documented years) of the branch. The age of the common ancestor was then reversely inferred using the mean CiMULE2 IL accumulation rate of the SWO phylogenetic clades in the homogeneous subset (not including primitive SWO) with the lowest amount of IL accumulations in Figure 4b right panel. TE family activation was inferred when the IL accumulate rate of the ancestor, or the mean IL accumulation rates of parallelly developed SWO lineages in case the ancestral IL accumulation rate is not available, is below 0.01 ILs/year, and the offspring had over 2-fold IL accumulation rate increase to ≥0.01 ILs/year. TE silencing was inferred when the offspring had less than half of ancestral IL accumulation rates.

### Identification of autonomous and non-autonomous transposons

The conserved DNA motifs/regions were searched in the terminal 200 bp regions of DVS transposon family members using MEME v5.3.0^55^. Peptides translated from all possible open reading frames from the CiMULE cluster representatives were output via the getorf tool from EMBOSS v6.6.0^56^. Conserved protein domains in the peptides were identified by searching (requiring E-value <1E-3) the Pfam 34.0 database^57^ using HMMER v3.3.2^58^. Transposon family members with conserved transposase domains were recognized as autonomous, and the rest were classified as non-autonomous.

### CiMULE2 transcription analysis

We downloaded 740 SWO, 171 pummelo, and 39 Mandarin RNA-seq datasets from NCBI (Table S12). The RNA-seq data were mapped to DVS with CiMULE2 members masked and a standalone member DVS_Mu2_1 (Figure S1). Transcriptomes including ≥ 20% rRNA reads were regarded as low-quality and excluded from further analysis, leaving 629 SWO (Table S13), 169 pummelo, and 39 mandarin transcriptomes useable. To minimize the methodological difference among the datasets, the total effective read count in each transcriptome was calculated as the number of mapped reads minus the rRNA reads. Then the normalized read abundance (FPKM) of CiMULE2 was calculated using the total number of reads uniquely mapped to CiMULE2 in the reference.

To detect CiMULE2 ILs with outer transcription starting sites in the 629 SWO transcriptomes, we scanned for the uniquely mapped read pairs overlapping both the CiMULE2 terminal 300 bp region (upstream of open reading frame) and the neighboring 1,000 bp external regions at the ILs, and a minimum of 2 read pairs were required to identify the expression of CiMULE2 from an IL in a transcriptome.

### Functional enrichment test

We performed functional enrichment of the IL-affected genes using g:Profiler^59^. The whole-genome genes in the unmasked regions of DVS were used as the reference set, and the putative IL-affected genes were the test set. Fisher’s exact test was applied in the statistical analysis, and the obtained *p* values were adjusted by Bonferroni correction.

## Acknowledgments

We acknowledge the authors of previous studies including Wang et al. (2017), Wu et al. (2014), Wang et al. (2021), Liang et al. (2020), Wu et al. (2018), Wu et al. (2021), and Zhu et al. (2022) for making the sequencing data used in this study publicly available.

## Author contributions

F.L. and B.W. conceived and designed this project. B.W. designed and implemented the computational pipeline. B.W. and Y.C. performed the analyses. Y.D. and F.G. helped with analysis. B.W., Y.C., F.G. and F.L. wrote the paper. All authors have read and approved the final version of this paper.

## Data and materials availability

The accession numbers of all used whole-genome NGS data and RNA-seq data are available in Tables S3 and S12. Genomic Pacbio CCS data of two Valencia sweet orange accessions sequenced in this study have been deposited in NCBI under BioProject ID PRJNA1076522. Scripts used in transposon insertion loci scanning are available at https://github.com/TheLuoFengLab/Transposon-insertion-locus-simulation-and-scanning.

## Supplementary Materials

**Supplementary Notes**

**Figure S1 to S6**

**Tables S1 to S16**

## References

1. Wu, G.A. et al. Sequencing of diverse mandarin, pummelo and orange genomes reveals complex history of admixture during citrus domestication. Nat. Biotechnol. 32, 656–662 (2014).

2. Wu, B. et al. A chromosome-level phased Citrus sinensis genome facilitates understanding Huanglongbing tolerance mechanisms at the allelic level in an irradiation induced mutant. bioRxiv, 2022.2002.2005.479263 (2022).

3. Webber, H.J., Batchelor, L.D. & Reuther, W. The Citrus Industry. (Univ. California Press, 1967).

4. Wang, L. et al. Somatic variations led to the selection of acidic and acidless orange cultivars. Nat. Plants (2021).

5. Butelli, E. et al. Retrotransposons control fruit-specific, cold-dependent accumulation of anthocyanins in blood oranges. Plant Cell 24, 1242–1255 (2012).

6. Wu, G.A. et al. Sequencing of diverse mandarin, pummelo and orange genomes reveals complex history of admixture during citrus domestication. Nature Biotechnology 32, 656–662 (2014).

7. Shamel, A.D. & Pomeroy, C.S. Bud mutations in horticultural crops. Journal of Heredity 27, 487–494 (1936).

8. Springer, N.M., Lisch, D. & Li, Q. Creating Order from Chaos: Epigenome Dynamics in Plants with Complex Genomes. Plant Cell 28, 314–325 (2016).

9. Foster, T.M. & Aranzana, M.J. Attention sports fans! The far-reaching contributions of bud sport mutants to horticulture and plant biology. Horticulture research 5, 44 (2018).

10. Baduel, P. et al. Genetic and environmental modulation of transposition shapes the evolutionary potential of Arabidopsis thaliana. Genome Biol 22, 138 (2021).

11. Negi, P., Rai, A.N. & Suprasanna, P. Moving through the Stressed Genome: Emerging Regulatory Roles for Transposons in Plant Stress Response. Front Plant Sci 7, 1448 (2016).

12. Raynes, Y., Wylie, C.S., Sniegowski, P.D. & Weinreich, D.M. Sign of selection on mutation rate modifiers depends on population size. Proceedings of the National Academy of Sciences 115, 3422–3427 (2018).

13. Liu, H. & Zhang, J. Yeast Spontaneous Mutation Rate and Spectrum Vary with Environment. Curr Biol 29, 1584–1591 e1583 (2019).

14. Xiong, Z., Yin, H., Wang, N., Han, G. & Gao, Y. Chromosome-level genome assembly of navel orange cv. Gannanzao (Citrus sinensis Osbeck cv. Gannanzao). G3 (Bethesda) 14 (2024).

15. Wang, L. et al. Somatic variations led to the selection of acidic and acidless orange cultivars. Nature Plants 7, 954–965 (2021).

16. Webber, H.J., Batchelor, L.D. & Reuther, W. History and development of the citrus industry. (Univ. California Press, 1967).

17. Duarte, A., Fernandes, J., Bernardes, J. & Miguel, G. Citrus As A Component Of The Mediterranean Diet. *Journal of Tourism*, Sustainability and Well-being 4, 289–304 (2016).

18. Tabara, H. et al. The rde-1 gene, RNA interference, and transposon silencing in C. elegans. Cell 99, 123–132 (1999).

19. Yang, J. et al. SWI3B and HDA6 interact and are required for transposon silencing in Arabidopsis. Plant J 102, 809–822 (2020).

20. Lanciano, S. & Cristofari, G. Measuring and interpreting transposable element expression. Nature reviews. Genetics 21, 721–736 (2020).

21. Zeilmaker, T. et al. DOWNY MILDEW RESISTANT 6 and DMR6-LIKE OXYGENASE 1 are partially redundant but distinct suppressors of immunity in Arabidopsis. Plant J 81, 210–222 (2015).

22. Aguilar-Martínez, J.A., Poza-Carrión, C. & Cubas, P. Arabidopsis BRANCHED1 acts as an integrator of branching signals within axillary buds. The Plant cell 19, 458–472 (2007).

23. Zhang, F. et al. Reprogramming of Stem Cell Activity to Convert Thorns into Branches. Current biology : CB 30, 2951–2961.e2955 (2020).

24. Ohnishi, T. et al. C-23 hydroxylation by Arabidopsis CYP90C1 and CYP90D1 reveals a novel shortcut in brassinosteroid biosynthesis. The Plant cell 18, 3275–3288 (2006).

25. Matus, J.T. et al. A group of grapevine MYBA transcription factors located in chromosome 14 control anthocyanin synthesis in vegetative organs with different specificities compared with the berry color locus. The Plant Journal 91, 220–236 (2017).

26. Sun, W. et al. Biochemical and Molecular Characterization of a Flavonoid 3-O-glycosyltransferase Responsible for Anthocyanins and Flavonols Biosynthesis in Freesia hybrida. Frontiers in Plant Science 7 (2016).

27. Alquezar, B., Rodrigo, M.J., Lado, J. & Zacarías, L. A comparative physiological and transcriptional study of carotenoid biosynthesis in white and red grapefruit (Citrus paradisi Macf.). Tree Genetics & Genomes 9, 1257–1269 (2013).

28. Griffiths, J. et al. Genetic characterization and functional analysis of the GID1 gibberellin receptors in Arabidopsis. The Plant cell 18, 3399–3414 (2006).

29. Gou, X. et al. Genetic evidence for an indispensable role of somatic embryogenesis receptor kinases in brassinosteroid signaling. PLoS genetics 8, e1002452 (2012).

30. Kidwell, M.G. & Lisch, D.R. Perspective: transposable elements, parasitic DNA, and genome evolution. Evolution 55, 1–24 (2001).

31. Fedoroff, N.V. Transposable Elements, Epigenetics, and Genome Evolution. Science 338, 758–767 (2012).

32. Malone, C.D. & Hannon, G.J. Molecular evolution of piRNA and transposon control pathways in Drosophila. Cold Spring Harb Symp Quant Biol 74, 225–234 (2009).

33. Senft, A.D. & Macfarlan, T.S. Transposable elements shape the evolution of mammalian development. Nat Rev Genet 22, 691–711 (2021).

34. Alseekh, S., Scossa, F. & Fernie, A.R. Mobile Transposable Elements Shape Plant Genome Diversity. Trends Plant Sci 25, 1062–1064 (2020).

35. Foster, T.M. & Aranzana, M.J. Attention sports fans! The far-reaching contributions of bud sport mutants to horticulture and plant biology. Horticulture research 5, 44 (2018).

36. Sun, X. et al. The role of transposon inverted repeats in balancing drought tolerance and yield-related traits in maize. Nat Biotechnol 41, 120–127 (2023).

37. Wang, L. et al. Somatic variations led to the selection of acidic and acidless orange cultivars. Nat Plants 7, 954–965 (2021).

38. Wu, B. et al. A chromosome-level phased genome enabling allele-level studies in sweet orange: a case study on citrus Huanglongbing tolerance. Horticulture Research (2022).

39. Foria, S. et al. Gene duplication and transposition of mobile elements drive evolution of the Rpv3 resistance locus in grapevine. Plant J 101, 529–542 (2020).

40. Wicker, T. et al. A unified classification system for eukaryotic transposable elements. Nat Rev Genet 8, 973–982 (2007).

41. Hickman, A.B. & Dyda, F. Mechanisms of DNA Transposition. Microbiol Spectr 3, MDNA3-0034-2014 (2015).

42. Belyayev, A. Bursts of transposable elements as an evolutionary driving force. Journal of Evolutionary Biology 27, 2573–2584 (2014).

43. Barrick, J.E. et al. Genome evolution and adaptation in a long-term experiment with Escherichia coli. Nature 461, 1243–1247 (2009).

44. Pal, C., Maciá, M.D., Oliver, A., Schachar, I. & Buckling, A. Coevolution with viruses drives the evolution of bacterial mutation rates. Nature 450, 1079–1081 (2007).

45. Gregory, T.R. Artificial Selection and Domestication: Modern Lessons from Darwin’s Enduring Analogy. Evolution: Education and Outreach 2, 5–27 (2009).

46. Seczynska, M., Bloor, S., Cuesta, S.M. & Lehner, P.J. Genome surveillance by HUSH-mediated silencing of intronless mobile elements. Nature (2021).

47. Slotkin, R.K. & Martienssen, R. Transposable elements and the epigenetic regulation of the genome. Nature reviews. Genetics 8, 272–285 (2007).

48. Li, H. Minimap2: pairwise alignment for nucleotide sequences. Bioinformatics 34, 3094– 3100 (2018).

49. Li, W. & Godzik, A. Cd-hit: a fast program for clustering and comparing large sets of protein or nucleotide sequences. Bioinformatics 22, 1658–1659 (2006).

50. Yan, H., Bombarely, A. & Li, S. DeepTE: a computational method for de novo classification of transposons with convolutional neural network. Bioinformatics 36, 4269–4275 (2020).

51. Larkin, M.A. et al. Clustal W and Clustal X version 2.0. *Bioinformatics (Oxford*, England) 23, 2947–2948 (2007).

52. Quinlan, A.R. & Hall, I.M. BEDTools: a flexible suite of utilities for comparing genomic features. Bioinformatics 26, 841–842 (2010).

53. Li, H. Aligning sequence reads, clone sequences and assembly contigs with BWA-MEM. arXiv preprint, arXiv:1303.3997v1302 (2013).

54. Li, H. et al. The Sequence Alignment/Map format and SAMtools. Bioinformatics 25, 2078–2079 (2009).

55. Bailey, T.L., Johnson, J., Grant, C.E. & Noble, W.S. The MEME Suite. Nucleic Acids Research 43, W39–49 (2015).

56. Rice, P., Longden, I. & Bleasby, A. EMBOSS: The European Molecular Biology Open Software Suite. Trends in Genetics 16, 276–277 (2000).

57. Mistry, J. et al. Pfam: The protein families database in 2021. Nucleic Acids Research 49, D412–D419 (2021).

58. Mistry, J., Finn, R.D., Eddy, S.R., Bateman, A. & Punta, M. Challenges in homology search: HMMER3 and convergent evolution of coiled-coil regions. Nucleic Acids Research 41, e121 (2013).

59. Raudvere, U. et al. g:Profiler: a web server for functional enrichment analysis and conversions of gene lists (2019 update). Nucleic Acids Research 47, W191–W198 (2019).

