## Supplementary Notes for "Transposon activities drive the selection and diversification of sweet orange (Citrus × sinensis) cultivars in the last 500 years"

**Transposon activity in sweet orange breeding**

**Supplementary Notes**

**Supplementary Note 1. Optimizations made for weet orange TE insertion scanning approach**

SUL1 and SUL1’ are the lengths of the shortest unique K-Mers at upstream and downstream of the ILs in the reference genome, namely, the shortest regions required to be covered for an alignment to be regarded as unique. Similarly, SUL2 is the length of the shortest unique K-Mer starting from the 5’ end of a TE member/group/family, and SUL2’ denotes the shortest unique K-Mer ending at its 3’ end. As shown in Figure 1c, to recall an IL, a read pair must contain two segments uniquely mapped to the upstream (or downstream) IL regions and the 5’ terminal (or 3’ terminal) of the scanned TE. While to precisely locate the coordinate of an IL, a read must cover both SUL1 and SUL2 or SUL1’ and SUL2’. Then, the shorter the SULs, the easier we can precisely locate an IL.

Under only one assumption that the sequencing reads are randomly distributed in the genome, we carried out in silico simulations of IL scanning to show the effects of the different parameters on IL recalling and locating. Under ≥ 300 bp insert size, ≤ 50 bp SUL (assuming a simplified model in which SUL1, 2, 1’, and 2’ all equal SUL), and a 100% mutant ratio (non-chimeric), the results show that a 5× sequencing depth achieves an average IL recalling rate of 99.0 ± 0.3% (Figure S5a). For ILs with 10% to 100% mutant ratios, under 150 bp read length and 50 bp SUL, a 15× sequencing depth can recall ILs from 97.3 ± 0.5% to 100.0 ± 0.0% (Figure S5b), and precisely locate them from 59.9 ± 1.7% to 99.3 ± 0.2% (Figure S5c). The precise coordinates of an IL would fail to be recalled when both (SUL1 + SUL2) and (SUL1’ + SUL2’) exceed the read length (Figure S5d).

The diploid SWO genome is highly heterozygous and contains a proportion of highly divergent regions (< 95% nucleotide similarity). Thus, to increase the IL recalling rates in divergent regions, we use the phased DVS genome as the reference, which has 98.5% completeness (estimated with 19 bp K-Mer), 25.6% higher than the haploid SWO genome used in the study of Wang et al. (2021)^1^. Because the allelically identical regions in the phased reference would increase the IL surrounding SULs, we masked the allelically identical regions in DVS_B of DVS (steps 1 and 2 in Figure 1b). First, the haploid chromosome set DVS_A (GCA_022201045.1) in DVS was split into 50 bp sliding windows with a step size of 10 bp; all window sequences were output and mapped to the other haploid chromosome set (DVS_B, GCA_022201065.1) via BWA v0.7.17. Then perfectly aligned (no mismatch, indel or clipped bases) regions in DVS_B were merged using BEDTools v2.29.2, and ≥150 bp merged regions were masked except for the two terminal 50 bp.

**Supplementary Note 2. Phylogeny of transposon families and terminal haplotype diversity**

For each TE family, DVS cluster representatives and all members from PTR (*C. trifoliata*)^2^, MSYJ (mandarin)^3^, and HWB (pummelo)^4^ assemblies were subjected to phylogenetic analysis by both Maximum Likelihood (ML) and Bayesian inference (BI). We carried out ML tree construction using IQ-TREE v2.1.3^5^ with automatic model selection and 1,000 bootstrap replicates. The BI method was applied using Mrbayes v3.2.7^6^ with the optimal model from IQ-TREE model selection, 110,000 generations of Markov chain Monte Carlo (MCMC) sampling, 1/100 tree sampling frequency and 100 burn-in trees. We visualized the phylogenetic tree using iTOL (Interactive Tree Of Life) v5^7^.

The multiple copies of identical or low-difference transposable elements in the reference genome will result in long SULs at their terminals. Therefore, we will not discover the ILs of these identical TEs due to the multiple alignments of their NGS reads/read segments. To reduce the SULs of TEs from several thousand bp to within 100 bp, we mask the duplicate copies and terminals of the transposable elements in the reference genome (Figure 1b). Here, CiMULE1 and CiMULE2 are used as examples. We first compared the haplotypes with the terminal 50 bp of CiMULE1 cluster members and assign them into 7 groups (MU1A to G) according to distinct terminal haplotypes at both ends (Figure S1a,b,c). All three CiMULE2 clusters only have one terminal 50 bp haplotype combination and are assigned into one group (Figure S1b,c,d,e). Furthermore, the neighboring bases of the TE members in the reference genome could interfere with the mapping of the reads from novel ILs. We mask all CiMULE1 and CiMULE2 members in the reference genome by the ambiguous base N, and add standalone copies of 17 CiMULE1 cluster representatives and 1 CiMULE2 (DVS_Mu2_1) (Figure 1b). To further distinguish the novel ILs of CiMULE1 clusters in MU1A and MU1E, we mask the 50 bp duplicate terminals of some of the cluster representatives. An example of the mapping of reads/read pairs crossing the IL boundary is shown in Figure 1d.

We compared the proportion of unique K-Mer with 20 to 100 bp length between the entire reference genome the surrounding sequences of the CiMULE ILs detected from the 11 SWO assemblies. The counts for K-Mers with lengths ranging from 20 to 100 bp in the entire modified reference genome and the SULs at CiMULE terminals and in the IL surrounding sequences were analyzed using our bash script based on Jellyfish v2.3.0^1^. In the modified reference genome, 36.8% 20 bp K-Mers, 72.0% 50 bp K-Mers, and 89.3% 100 bp K-Mers are unique. The shortest unique K-Mers of both CiMULE1 and CiMULE2 ILs in the 11 SWO long-read assemblies are significantly enriched (p < 0.05 by Chi-squared test) in those with lengths ≤ 45 bp (Figure S6a,b). Only 4 of the 69 CiMULE1 ILs have both SULs > 100bp, 3 of which fail to be scanned. No CiMULE2 IL has both SULs > 100bp. The terminal SULs of the CiMULE1 representatives in the modified reference range from 20 to 71 bp (Figure S6c), averaged at 33.3 bp. The CiMULE2 representative (DVS_Mu2_1) in the modified reference genome has a 32 bp SUL2 and a 59 bp SUL2’. These statistical data implied our pipeline could achieve high recall rates in sweet orange. Without making these optimizations, only 267 mutator insertions with <90% accuracy were detected in 114 SWO accessions, while we have observed 751 CiMULE1 and 2,596 CiMULE2 ILs with high accuracy (>99% with PCR-free sequencing data as shown below) in the same dataset.

**Supplementary Note 3. Insertion locus loss comparison among different TE types**

Transposable elements have different transposition mechanisms. Retrotransposons acquire new insertions in the genome in a copy and paste manner. Most DNA transposons use the cut and paste method except for the helitrons that transpose without excisions. Both transposition with and without excision have been observed for MULEs, and mechanisms of duplication without excisions were only hypothesized to exist in the germinal tissues^8^.

Theoretically, transpositions with and without excisions will have different IL loss rates in a population. We compared the parental IL loss ratios of the different TE types in SWO. We inferred ILs present in at least three of the four primitive SWO accessions and both mandarin and pummelo to be SWO parental ILs, which should be detected in any SWO accession if there had been no excision. To minimize the impact of IL recalling difference, we only used ILs with both SULs no longer than 55 bp in IL loss analysis, which would assure ~99% IL recall rates in all the SWO accessions in simulation. We observed significantly (FDR<0.01) lower IL loss ratio on the retrotransposon type (LTR) than three (CACTA, Harbinger, and hAT) of the four DNA transposon types (Figure 1f). The last DNA transposon superfamily MULE has significantly (FDR<0.01) lower IL loss rate than the other three, but no statistical difference (p=0.92) from LTR in SWO. The IL loss rates of the five different types of TEs are limited, ranging from 1.6% (MULE) to 4.0% (CACTA), which should produce limited noise on IL-based phylogenetic analysis.

**Supplementary Note 4**. **More transposon insertion loci putatively contributing to sweet orange horticultural traits**

Two CiMULE2 ILs in intronic and 5’-UTR regions of the same gene *CsGAI,* a repressor of the gibberellin (GA) signaling pathway, were detected in MIDNIT and LJG, respectively, which may explain why MIDNIT grows more slowly and ripens earlier than other Valencia oranges^9^. Two other CiMULE2 ILs are found in intronic and 5’-UTR regions of a *CsKAO2* allele in four Navel oranges and a seedless Jincheng, respectively, and *CsKAO2* catalyzes the key step in the biosynthesis of GAs^10^.

In Bingtangcheng, nine genes are putatively affected by eight CiMULE2 tag ILs. Two tag ILs are located upstream of *CsMUR3* and *CsLBD19* and putatively affect the salt tolerance^11^ and organ morphogenesis^12^, respectively.

Seven Jincheng tag CiMULE2 ILs putatively affect seven genes. A tag CiMULE2 IL is located upstream of *CsGHR1*, which induces stomatal movement through regulation of anion channel activity in response to ABA, H_2_O_2_, diurnal light/dark transitions, and high CO_2_ concentration. This IL putatively has an impact on its fruit and seed development, since mature citrus fruit generally has closed stomata which prevents water loss, but the closure of stomata in the early development stage of citrus fruit reduces transpiration and disrupts seed development^13^.

In Valencia orange, we find three tag CiMULE2 ILs upstream of *CsGID1* & *CsPMT22*, *CsSERK1*, and a C2H2-type zinc finger family gene. *CsGID1* is an ortholog of the Arabidopsis *AtGID1* family, which encode soluble GA receptors playing key roles in GA signaling and is involved in multiple development processes^14^ including ovary development^15^. GA plays an important role in citrus fruit development including fruit set^16^ and fruit maturation^17^. *CsSERK1* is an ortholog of both *AtSERK1* and *AtSERK2*, which play an indispensable role in BR signaling pathway in *Arabidopsis^18^* and control male sporogenesis^19^. Moreover, BR treatment on citrus has been reported to enhance its biotic^20^ and abiotic^21^ stress tolerance, and promote fruit setting^22^.

Navel orange possesses three CiMULE1 and two CiMULE2 tag ILs. A tag CiMULE1 IL is located at upstream of a *CYP* family member (DVS9B02337), and *CYP* family members are related with important roles in plant development and defense^23^.

**References**

1	Wang, L. *et al.* Somatic variations led to the selection of acidic and acidless orange cultivars. *Nature Plants* **7**, 954–965 (2021). <https://doi.org:10.1038/s41477-021-00941-x>

2	Peng, Z. *et al.* A chromosome‐scale reference genome of trifoliate orange ( Poncirus trifoliata ) provides insights into disease resistance, cold tolerance and genome evolution in Citrus. *The Plant Journal* **104**, 1215-1232 (2020). <https://doi.org:10.1111/tpj.14993>

3	Wang, L. *et al.* Genome of wild mandarin and domestication history of mandarin. *Molecular plant* **11**, 1024–1037 (2018). <https://doi.org:10.1016/j.molp.2018.06.001>

4	Wang, X. *et al.* Genomic analyses of primitive, wild and cultivated citrus provide insights into asexual reproduction. *Nature Genetics* **49**, 765–772 (2017). <https://doi.org:10.1038/ng.3839>

5	Minh, B. Q. *et al.* IQ-TREE 2: New Models and Efficient Methods for Phylogenetic Inference in the Genomic Era. *Molecular biology and evolution* **37**, 1530–1534 (2020). <https://doi.org:10.1093/molbev/msaa015>

6	Ronquist, F. *et al.* MrBayes 3.2: efficient Bayesian phylogenetic inference and model choice across a large model space. *Systematic biology* **61**, 539–542 (2012). <https://doi.org:10.1093/sysbio/sys029>

7	Letunic, I. & Bork, P. Interactive Tree Of Life (iTOL) v5: an online tool for phylogenetic tree display and annotation. *Nucleic Acids Research* **49**, W293-W296 (2021). <https://doi.org:10.1093/nar/gkab301>

8	Lisch, D. Mutator transposons. *Trends in Plant Science* **7**, 498–504 (2002). <https://doi.org:10.1016/s1360-1385(02)02347-6>

9	Webber, H. J., Batchelor, L. D. & Reuther, W. *The Citrus Industry*. 1-39 (Univ. California Press, 1967).

10	Helliwell, C. A., Chandler, P. M., Poole, A., Dennis, E. S. & Peacock, W. J. The CYP88A cytochrome P450, ent-kaurenoic acid oxidase, catalyzes three steps of the gibberellin biosynthesis pathway. *Proceedings of the National Academy of Sciences* **98**, 2065–2070 (2001). <https://doi.org:10.1073/pnas.98.4.2065>

11	Li, W., Guan, Q., Wang, Z.-Y., Wang, Y. & Zhu, J. A bi-functional xyloglucan galactosyltransferase is an indispensable salt stress tolerance determinant in Arabidopsis. *Molecular Plant* **6**, 1344–1354 (2013). <https://doi.org:10.1093/mp/sst062>

12	Matsumura, Y., Iwakawa, H., Machida, Y. & Machida, C. Characterization of genes in the ASYMMETRIC LEAVES2/LATERAL ORGAN BOUNDARIES (AS2/LOB) family in Arabidopsis thaliana, and functional and molecular comparisons between AS2 and other family members. *The Plant journal : for cell and molecular biology* **58**, 525–537 (2009). <https://doi.org:10.1111/j.1365-313X.2009.03797.x>

13	Lugassi, N. *et al.* Expression of Hexokinase in Stomata of Citrus Fruit Reduces Fruit Transpiration and Affects Seed Development. *Frontiers in plant science* **11**, 255 (2020). <https://doi.org:10.3389/fpls.2020.00255>

14	Griffiths, J. *et al.* Genetic characterization and functional analysis of the GID1 gibberellin receptors in Arabidopsis. *The Plant cell* **18**, 3399–3414 (2006). <https://doi.org:10.1105/tpc.106.047415>

15	Ferreira, L. G. *et al.* GID1 expression is associated with ovule development of sexual and apomictic plants. *Plant cell reports* **37**, 293–306 (2018). <https://doi.org:10.1007/s00299-017-2230-0>

16	Bermejo, A. *et al.* Auxin and Gibberellin Interact in Citrus Fruit Set. *Journal of Plant Growth Regulation* **37**, 491–501 (2018). <https://doi.org:10.1007/s00344-017-9748-9>

17	Alferez, F., Carvalho, D. U. d. & Boakye, D. Interplay between Abscisic Acid and Gibberellins, as Related to Ethylene and Sugars, in Regulating Maturation of Non-Climacteric Fruit. *International journal of molecular sciences* **22** (2021). <https://doi.org:10.3390/ijms22020669>

18	Gou, X. *et al.* Genetic evidence for an indispensable role of somatic embryogenesis receptor kinases in brassinosteroid signaling. *PLoS genetics* **8**, e1002452 (2012). <https://doi.org:10.1371/journal.pgen.1002452>

19	Albrecht, C., Russinova, E., Hecht, V., Baaijens, E. & Vries, S. d. The Arabidopsis thaliana SOMATIC EMBRYOGENESIS RECEPTOR-LIKE KINASES1 and 2 control male sporogenesis. *The Plant cell* **17**, 3337–3349 (2005). <https://doi.org:10.1105/tpc.105.036814>

20	Canales, E. *et al.* 'Candidatus Liberibacter asiaticus', causal agent of citrus Huanglongbing, is reduced by treatment with brassinosteroids. *PloS one* **11**, e0146223 (2016). <https://doi.org:10.1371/journal.pone.0146223>

21	Ghorbani, B. & Pakkish, Z. Brassinosteroid enhances cold stress tolerance of Washington navel orange (Citrus sinensis L.) fruit by regulating antioxidant enzymes during storage. *1331-7768* **79**, 109–114 (2014).

22	Sugiyama, K. & Kuraishi, S. Stimulation of fruit set of 'Morita' navel orange with brassinolide. *Acta Horticulturae*, 345–348 (1989). <https://doi.org:10.17660/ActaHortic.1989.239.54>

23	XU, J., WANG, X.-y. & GUO, W.-z. The cytochrome P450 superfamily: Key players in plant development and defense. *Journal of Integrative Agriculture* **14**, 1673–1686 (2015). <https://doi.org:10.1016/s2095-3119(14)60980-1>
