## Supplementary Figures for "Transposon activities drive the selection and diversification of sweet orange (Citrus × sinensis) cultivars in the last 500 years"

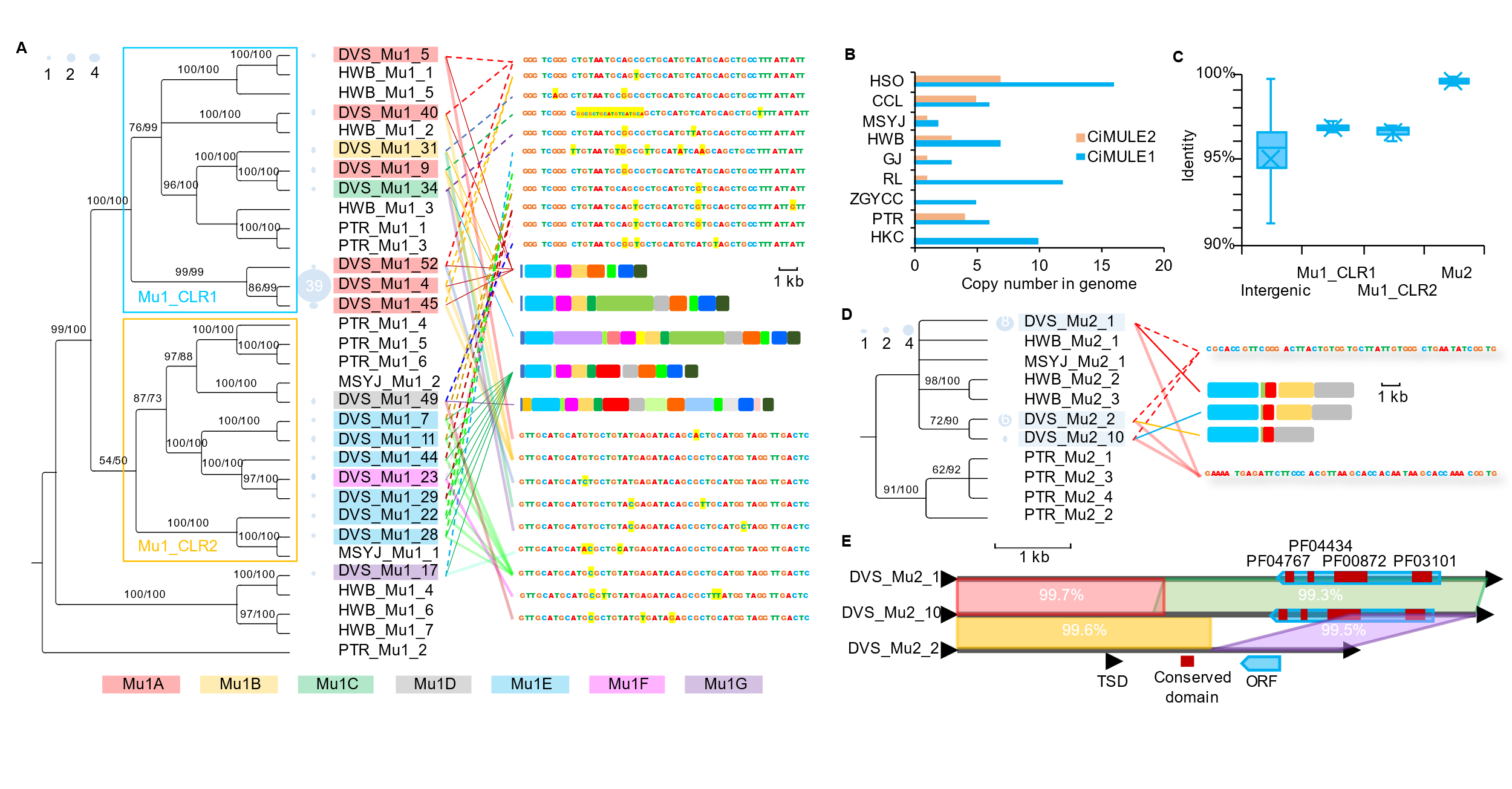

**Figure S1. Diversity of *Citrus* CiMULE1 and CiMULE2 transposon families. a** and **d**, The midpoint rooted phylogenetic trees (left), structural variants (middle right), and 50 bp terminal haplotypes (top and bottom right) of CiMULE1 and CiMULE2, respectively. CiMULE1 and CiMULE2 members from genomes of PTR (*Citrus trifoliata*), MSYJ (*Citrus reticulata*), and HWB (*Citrus maxima*), and 17 CiMULE1 and 3 CiMULE2 cluster representatives from the diploid Valencia sweet orange (DVS) genome were applied in the phylogenetic analyses. The bubbles left to the node names show the sizes of the corresponding clusters in DVS. Only branches supported by > 50 percent bootstrap (the number before slashes) tests using the Maximum likelihood method and with > 50 percent probabilities (after slashes) by Bayesian inference are shown in the phylogenetic trees. For the structural variant diagrams of either mutator family, the bins of the same colors denote homologous regions. The two CiMULE1 clusters (Mu1_CLR1 and Mu2_CLR2) are indicated by the blue and yellow frames. The CiMULE1 members from DVS are assigned into seven subgroups (Mu1A-G, highlighted with distinct background colors) sharing no terminal haplotype for insertion locus scanning. In the terminal haplotypes, minor alleles of the variants are highlighted with yellow backgrounds. **b**, Copy numbers of CiMULE1 and CiMULE2 members in the genomes of 8 *Citrus* species and *Atalantia buxfoliata*. HSO, di-haploid *Citrus × sinensis*; CCL, *Citrus × clementina*; GJ, *Citrus japonica*; RL, *Citrus medica*; ZGYCC, *Citrus ichangensis*; HKC, *Atalantia buxfoliata*. **c**, Boxplots showing the nucleotide identity distributions among PTR and DVS CiMULE1 and CiMULE2 members and other intergenic orthologous regions. The center line, median; fork, mean; boxes, first and third quartiles; whiskers, 5th and 95th percentiles; dots, outliers. **e**, Alignment and conserved protein domains of CiMULE2 cluster representatives from DVS. Homologous regions between the members are connected with colorful parallelograms with nucleotide identities shown inside. TSD, target site duplication; ORF, open reading frame. Conserved protein domains (from the Pfam 34.0 database): PF04767, Pox_F17, DNA-binding phosphoprotein; PF04434, SWIM zinc finger; PF00872, Mutator family transposase; PF03101, FAR1 DNA-binding domain.

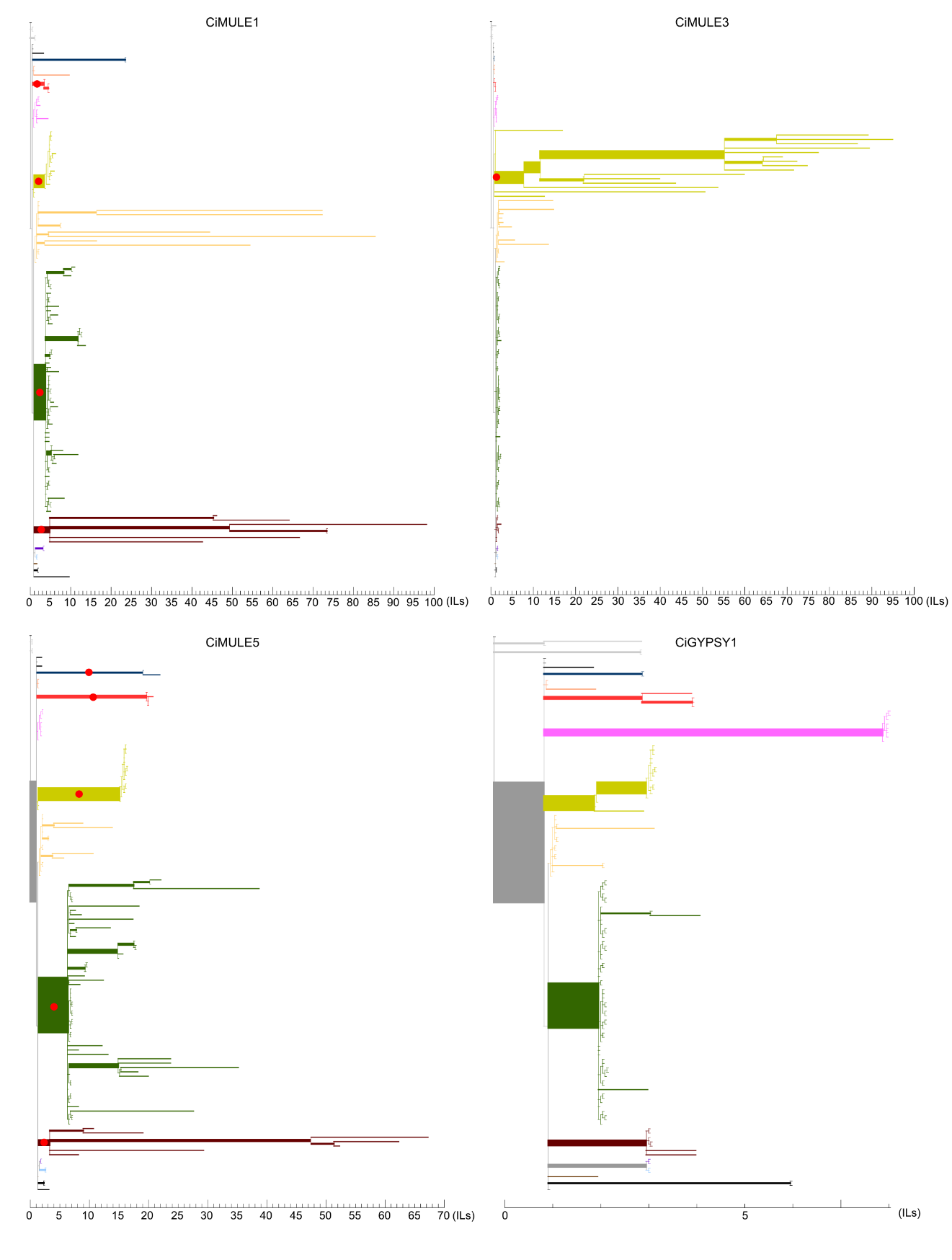

**Figure S2. Insertion locus (IL) accumulation of four boosted transposable element families on phylogenetic paths of sweet oranges.**

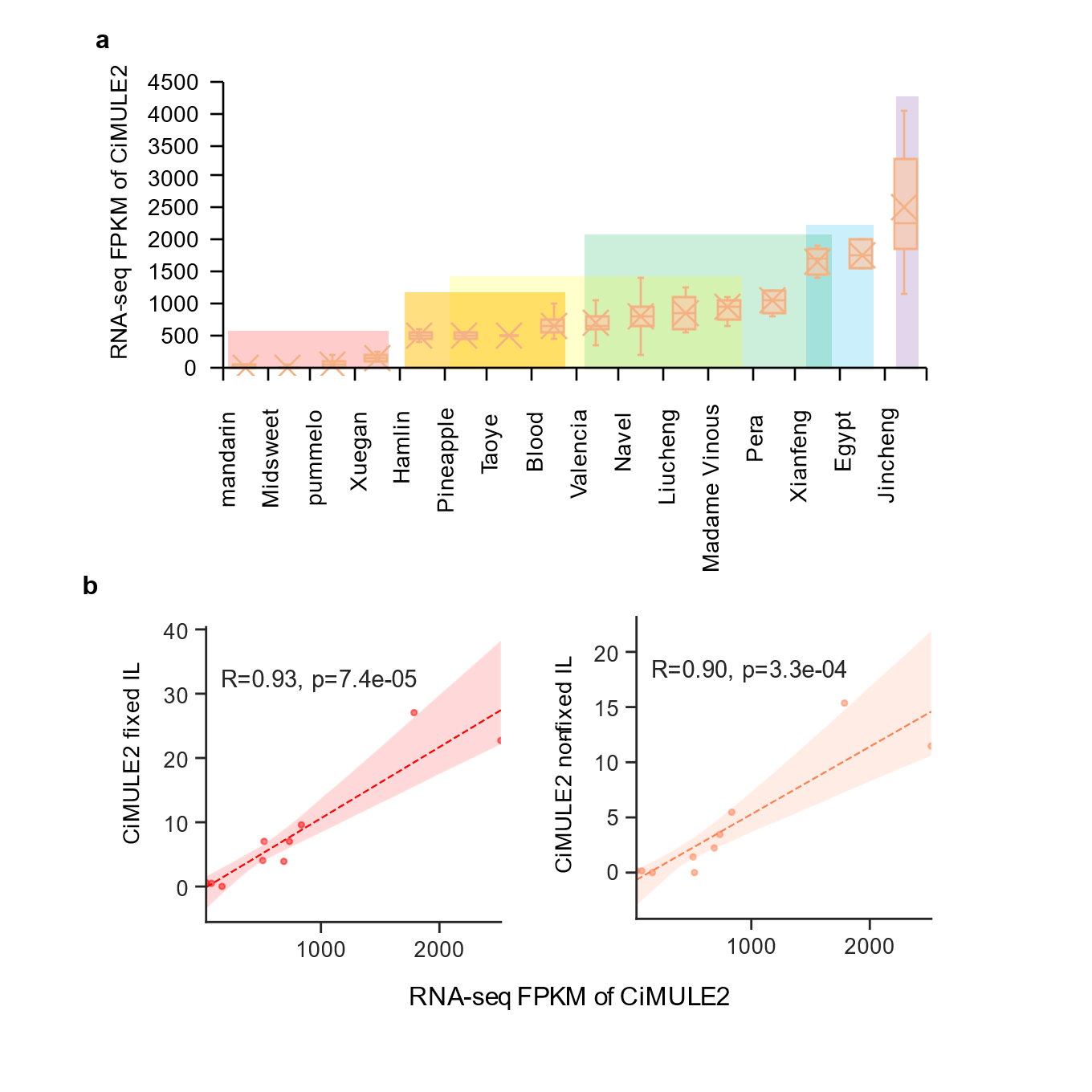

**Figure S3. Expression abundance of CiMULE2 in citrus transcriptomes and its correlation with genomic insertion locus counts. a**, Normalized RNA-seq abundance of CiMULE2 in mandarin, pummelo, and multiple sweet orange cultivar groups. FPKM, fragments per kilobase of transcript per million mapped reads. Two primitive SWO accessions, WHXG and WSXG, belong to the Xuegan group, and another primitive SWO accession TYC from the Taoye orange groups. In the boxplots: the center line, median; fork, mean; boxes, first and third quartiles; whiskers, 5th and 95th percentiles. **b**, Correlations between the mean RNA-seq abundance and the mean fixed (left) and non-fixed (right) CiMULE2 IL counts in mandarin, pummelo, and eight SWO cultivar groups. Fixed and non-fixed ILs had ≥ and < 90% mutant ratios in the owner accessions, respectively. The eight groups include Xuegan, Hamlin, Taoye, Blood, Valencia, Navel, Egypt, and Jincheng.

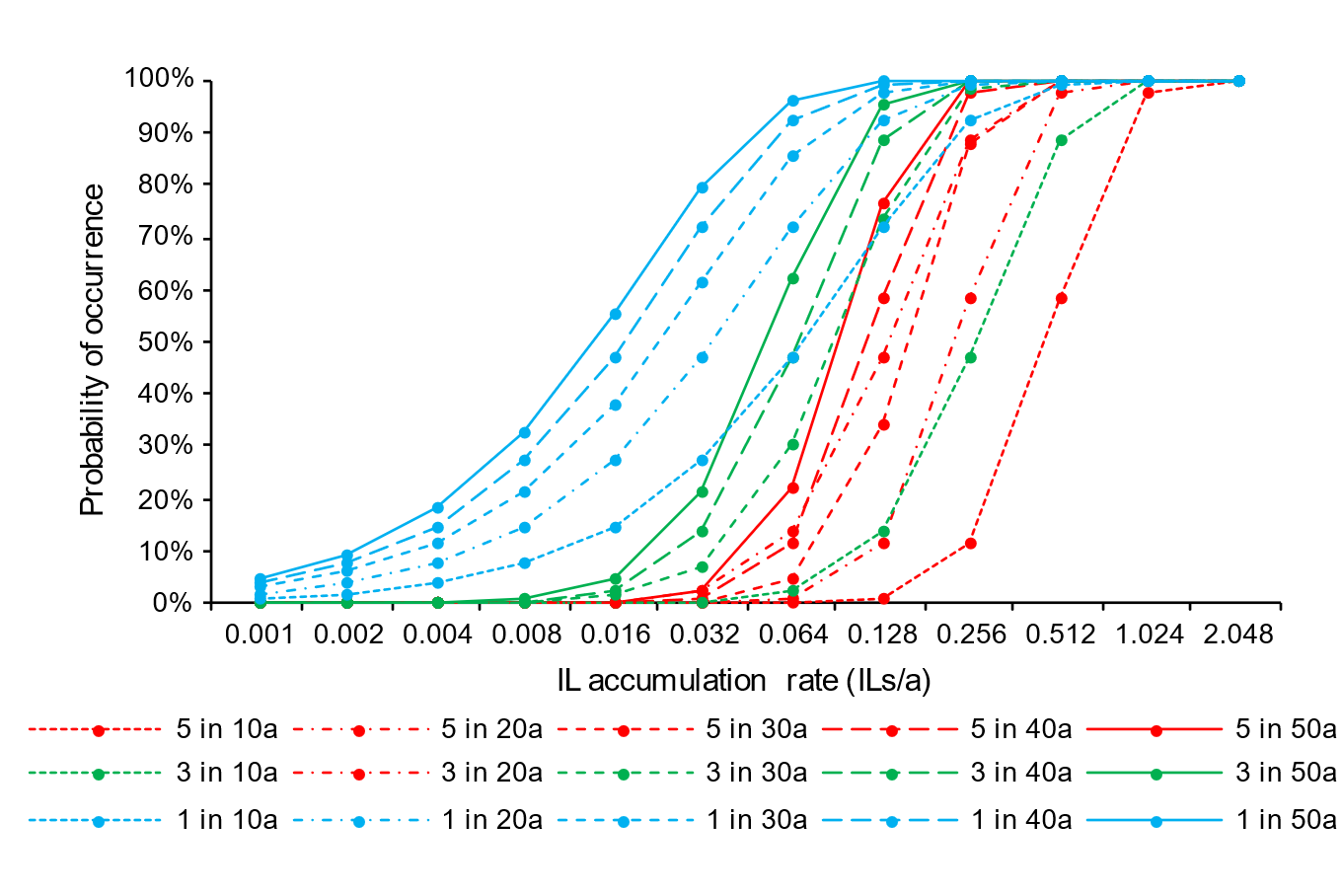

**Figure S4. Minimum requirements of transposon insertion locus (IL) accumulation rates for cultivar formation within decades.** The solid dots denote the probabilities of occurrence (vertical axis) of 1, 3, and 5 random ILs in 10 to 50 years (a) on a branch with 0.001 to 2.048 ILs/a accumulation rates. The formation of many sweet orange cultivars involved multiple functional ILs, which are only expected to account for a small proportion (estimated to be <0.01%) of naturally occurring ILs. Thus, the occurrence of ILs equal to the required functional IL count is only a minimum requirement.

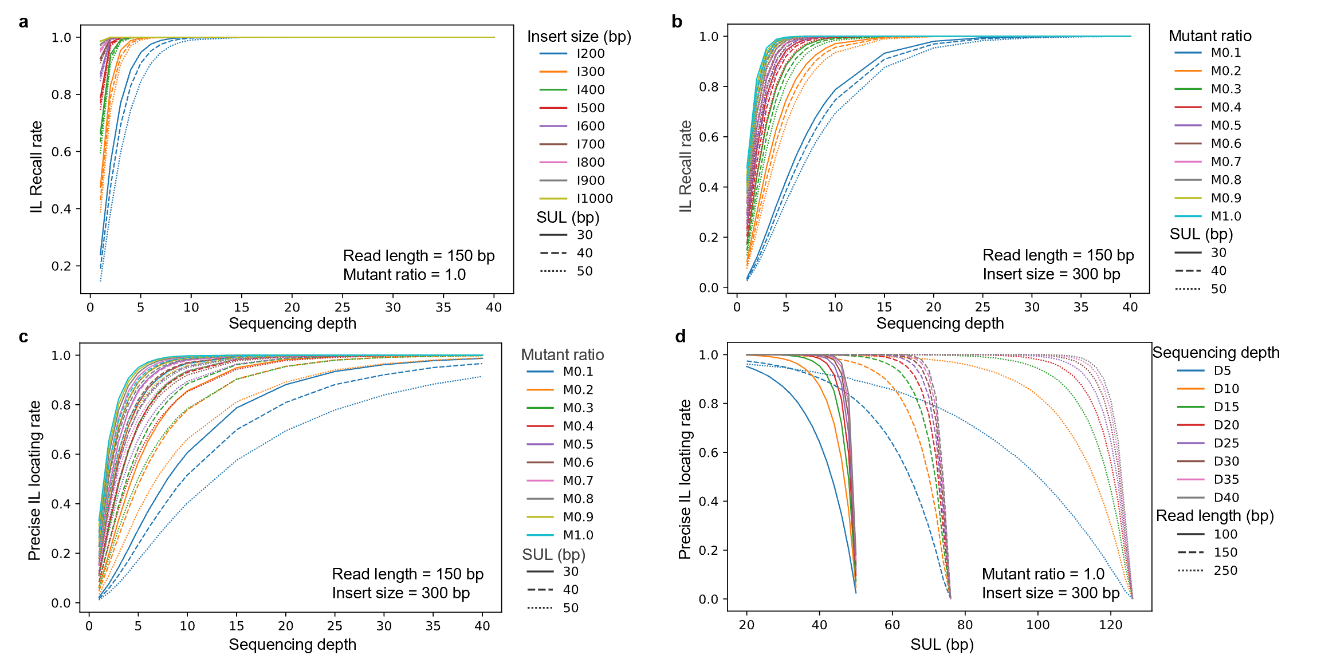

**Figure S5. Technical explanation and simulation of transposon insertion locus (IL) scanning.** SUL, the shortest unique K-Mer length. In panels A and B, the blue and green bars denote upstream and downstream regions of transposon ILs in the chromosomes and the deep red bars indicate the transposons. **a**, Mapping of pair-end sequencing reads to transposons embedded in the reference genome. IL overlapping read pairs are classified as informative and uninformative in recalling the IL. Precisely locating reads that could recall and precisely locate an IL, and recall-only read pairs could only recall an IL and locate it in a certain range. The leftmost and rightmost read pairs allowing recalling of the IL are shown at the bottom. The bottom lines of the green and purple triangles indicate the ranges of the first aligned bases on the reference for informative read pairs and reads capable of precisely locating the IL, respectively. **b**, Mapping of pair-end reads to novel chimeric ILs absent in the reference. Wild-type and mutant-type read pairs are drawn over and below the chromosome, respectively. Mutant-type read pairs are either dis-concordantly mapped or include at least one read split-mapped to both the IL surrounding sequences and the transposon. The segments or reads from the same mutant-type read pairs are marked with the same numeric ID. When calculating the mutant ratio, the effective wild-type read count (C_w_) is used to compensate for the uncounted IL uninformative read pairs. **c**, **d**, **e,** and **f**, *In silico* simulations of transposon IL recalling and precise locating with different parameter combinations. The model was simplified in simulations by assuming (SUL1 + SUL2) equal to (SUL1’ + SUL2’) so that only a single parameter SUL (the average of SUL1, SUL2, SUL1’ and SUL2’) was required in simulation. The average recall rates on 1,000 ILs × 100 iterations simulation are shown for each parameter combination in the graphs. Constant parameters applied in the simulation are shown in the bottom right of the four graphs.

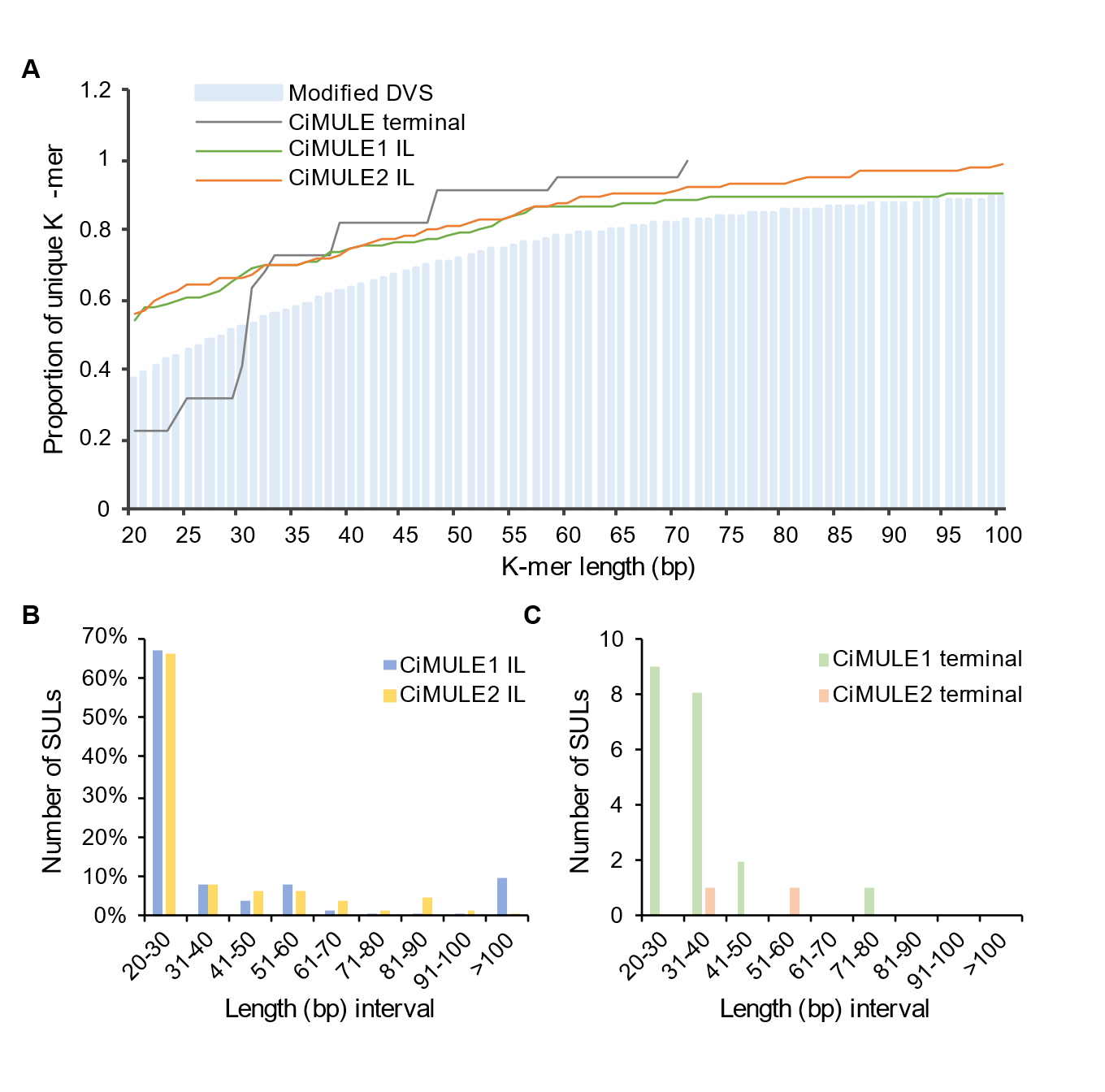

**Figure S6. Statistics of unique K-Mers in the reference, CiMULE1 and CiMULE2 terminals and their surrounding sequences.** All the unique K-Mers and shortest unique K-Mer lengths (SULs) were analyzed using the modified diploid Valencia sweet orange (DVS) genome (including the CiMULE1 and CiMULE2 members) applied in transposable element scanning. The CiMULE1 and CiMULE2 insertion loci (ILs) in the 11 sweet orange assemblies were used to estimate the unique K-Mer and SULs in the IL surrounding sequences. **a**, Proportions of unique K-Mers at given lengths under different circumstances. The light blue histogram shows the proportions of unique K-Mers at the corresponding K-Mer lengths in the modified DVS genome. The 20 distinct CiMULE1 terminals and 2 CiMULE2 terminals in the modified DVS genome are summarized together as the CiMULE terminal. The unique K-Mer ratios of the insertion locus (IL) surrounding sequences of CiMULE1 and CiMULE2 are shown as CiMULE1 IL and CiMULE2 IL in the legend. **b** and **c**, Distribution of the shortest unique K-Mer lengths (SULs) surrounding the CiMULE ILs and at their terminals.
